# Synthesis of Ultra-Large Fibrous Proteins from Bacteria via a Looped-Translation System

**DOI:** 10.64898/2026.07.16.738376

**Authors:** Qi Xie, Louis J. Papa, Anton M. Barybin, Michael Xiong, Matthew D. Shoulders, Stephen D. Fried

**Affiliations:** Department of Chemistry, Johns Hopkins University, Baltimore, MD 21218 USA; Department of Chemistry, Massachusetts Institute of Technology, Cambridge, MA 02139 USA; Koch Institute for Integrative Cancer Research, Massachusetts Institute of Technology, Cambridge, MA 02139 USA; Institute for Medical Engineering and Science, Massachusetts Institute of Technology, Cambridge, MA 02139 USA; Broad Institute of Massachusetts Institute of Technology and Harvard University, Cambridge, MA 02142; T. C. Jenkins Department of Biophysics, Johns Hopkins University, Baltimore, MD 21218 USA

## Abstract

High molecular weight fibrous proteins such as silk, elastin, and collagens, are fundamental for providing shape to macroscopic biological structures, yet their recombinant production remains challenging because of their extreme size and sequence repetitiveness. Here, we report a circular RNA-based ribosome translation platform that enables iterative ribosome synthesis of fibrous proteins through continuously “looped” translation. To promote efficient circularization of repetitive fibrous protein transcripts, we combined a synonymous codon locker sequence strategy with RNA circularization chaperones. Guided by a ribosome traffic model, we further optimized the translation bottlenecks within the circular RNA, substantially improving translation yields. The established looped translation platform is applicable to at least six classes of fibrous proteins and generated products with molecular weight exceeding titin at 3.8 MDa. The synthesized polypeptides were characterized through electron microscopy, bulk material fabrication, and mechanical analysis, demonstrating properties associated with ultra-high molecular weight polypeptides. Finally, we coupled looped translation to secretion through a programmed ribosomal frameshift, enabling export of fibrous protein across cellular membranes in both *Escherichia coli* and *Bacillus subtilis*. We envision that the genetic tools presented here could find a range of applications in bioplastics and engineered living materials.

## Main

Ultra-high molecular weight polymers, such as ultra-high molecular weight polyethylene (UHMWPE), exhibit unique mechanical properties due to their ability to transfer loads through extensive covalent networks^1,2^. In nature, proteins perform a wide variety of functions such as self-assembly, molecular sensing, catalysis, and much more owing to the complexity and versatility of the twenty canonical amino acids. However, there are very few examples of natural polypeptides with ultra-high molecular weight scale (99% of proteins in the Uniprot database are <1500 amino acids)^3^. Despite their rarity, natural mega-proteins exhibit fascinating properties facilitated by their massive size. For instance, spider silk proteins form fibers with exceptional toughness^4,5^, while titin, the largest protein in the human proteome (ca. 27,000–35,000 amino acids), functions as a molecular spring that provides passive elasticity to muscle^6^. Hence, for certain mechanical functions, protein sequences were selected to expand to ultra-high chain lengths rather than through multivalency and self-assembly^7,8^.

To replicate these structures and functions, synthetic biologists have attempted various strategies to encode and express high molecular weight proteins. DNA-level approaches, such as head-to-tail ligation^4^ and rolling circle amplification^9^, have been used to create tandem repeat genes of silk proteins^10^, elastin-like polypeptides^11^, and squid ring tooth proteins up to 285 kDa. Because highly repetitive sequences are often prone to instability from recombinase activity and difficult to propagate, computational approaches for codon optimization and sequence scrambling have been deployed to reduce sequence repetitiveness for a range of genes encoding repetitive polypeptides, including elastin-like and resilin-like proteins^12^. Orthogonal sets of protein-centric strategies focus on functionalizing the termini of a unit polypeptide with reactive motifs that enable intermolecular ligation, thereby mimicking classical polymerization reactions. Reactive domains such as split inteins^13–15^, the spycatcher/spytag system^16^, and sortase/sortase recognition motif ^17^ have all been explored in this context.

Circular mRNAs, which allow ribosomes to iteratively translate an open reading frame indefinitely^18^, represent another strategy to produce highly repetitive ultra-high molecular weight proteins that can avoid some of the pitfalls of DNA-based and protein-centric approaches. Namely, DNA-based approaches are limited by the size of the gene needed to encode a mega-protein, and protein-centric approaches require many non-encoded, non-templated ligation events to occur, which limits the programmability of the polypeptide sequence generated. Synthetic circular mRNAs can be generated using chemical^19^ and enzymatic ligation^20^, or through ribozyme-based methods amenable to circularize RNA both *in vitro* and *in vivo*. In particular, group I self-splicing introns, initially discovered in *Tetrahymena thermophila*^21^, can be engineered in a permuted-intron-exon structure (Fig. 1a)^22^, and have been successfully executed using the introns from T4 phage thymidylate synthase (td) and *Anabaena* pre-tRNA genes.

**Fig. 1.**
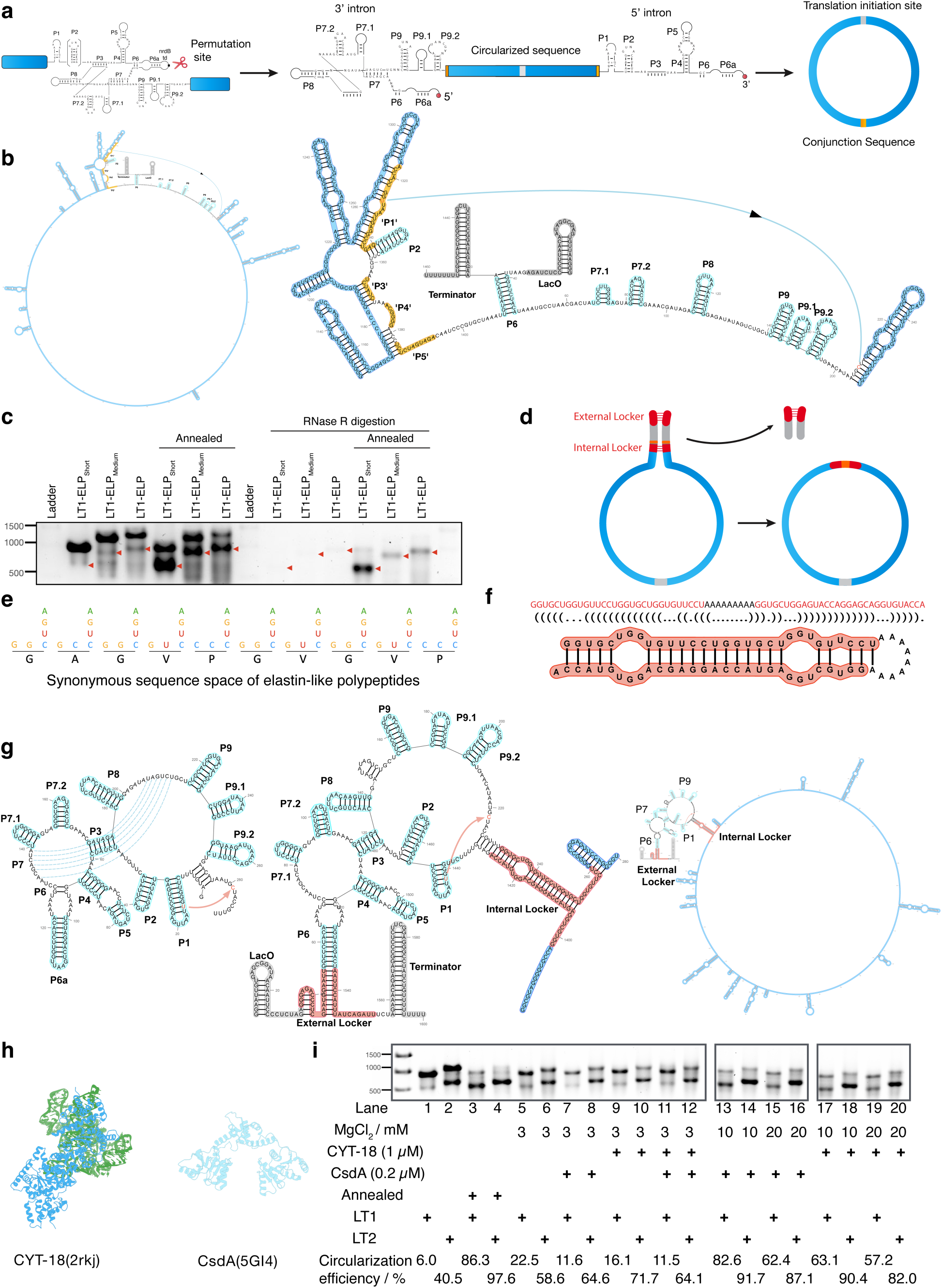
Engineering RNA circularization of fibrous protein coding sequences. a, Schematic of the permuted intron–exon (PIE) strategy for RNA circularization. The coding sequence is inserted between split intron fragments, enabling self-splicing and ligation into a circular RNA with a defined junction sequence. **b,** RNAfold prediction of LT1 precursor RNA. The global secondary structure is shown on the left, with zoomed-in views of intron regions on the right. T4 intron elements are colored cyan when correctly paired and yellow when mispaired. Coding sequences are shown in blue, while LacO and terminator sequences are shown in grey. **c,** Electrophoretic mobility shift assays and RNase R assays on *in vitro* transcribed/circularized RNA for constructs of varying length and with/without re-annealing. Red arrows denote the circularized product. **d,** Locker sequence strategy to promote intron folding and RNA circularization. **e,** Synonymous codon strategy to design the internal locker sequence based on the coding sequence space. **f,** Example of a synonymous internal locker sequence and its secondary structure. **g,** RNAfold prediction of circular RNA construct with synonymous locker sequences. The left panel shows the secondary structure of the T4 intron sequence. The middle panel shows the zoom-in view of intron sequences in the context of the circular mRNA constructs. The right panel shows the overview of RNAfold prediction for the whole circular RNA construct. **h,** Structure of protein factors that promote RNA circularization. **i,** In vitro RNA circularization assay for circular RNA constructs with synonymous locker sequences (LT2) and/or protein factors added *in trans*.

Perriman and Ares first demonstrated the permuted intron strategy to produce long GFP concatemers in *E. coli*^23^. Since that report, a number of studies have applied circular mRNA to produce concatemer proteins^24–26^, however, the technology has not been broadly applied beyond proof-of-concept or to protein materials to the best of our knowledge. Our own incipient attempts at circularizing mRNA encoding silk made us consider the possibility that sequences encoding fibrous proteins may be particularly challenging to circularize because they tend to be GC-rich^18^. Moreover, intracellular expression of large fibrous protein assemblies tends to be toxic. Recognizing these issues, we sought strategies to couple translation of circular mRNA to co-translational protein secretion so that growing polypeptides would be exported concomitant with their synthesis, thereby alleviating intracellular burden and promoting extracellular assembly.

Here, we incorporate design elements into permuted introns to render efficient circularization sequence-agnostic, including for repetitive GC-rich sequences common to fibrous proteins. We also exploit synonymous codon-locking architectures to tune RNA structural compatibility independently of the encoded protein sequence. In parallel, we deploy RNA trans-acting factors, including RNA chaperones and helicases, to actively remodel structured RNA intermediates and promote productive circularization of repetitive transcripts. Guided by a ribosome traffic model, we further optimized looped translation by minimizing sequence features predicted to slow translation during iterative ribosome cycling on circular RNAs. We moreover demonstrate the incorporation of a programmed ribosomal frameshift to ensure that looped translation can be coupled to vectorial translation of a signal sequence, enabling proper engagement with the bacterial translocon and therefore co-translational protein export of ultra-high molecular weight polypeptides.

### Circularizing mRNA Encoding Fibrous Protein Sequences Through Synonymous Locker Sequences and RNA Chaperones

When the group I self-splicing intron is circularly permuted at P6a, the engineered topology redirects the phosphoryl-group transfer reaction catalyzed by this ribozyme to circularize a segment of RNA in between the two half-introns, instead of splicing two exons together (Fig. 1a). Pioneering work by Perriman and Ares^23^ first demonstrated the capacity of group I introns to circularize mRNA in vivo, and subsequent work by us^18^ and others^25,27^ optimized this system to direct ribosomes to synthesize large repetitive proteins by looping translation. Our early attempts to utilize this system to circularize mRNAs encoding the elastin-like polypeptide (ELP, (G[A/V]GVP)_n_, a model fibrous protein^12,28–30^) were unsuccessful, however (Supplementary Fig. 1a-b)^18^. Fibrous protein sequences are generally highly repetitive and GC-rich, which we hypothesized could preclude the proper folding and assembly of the catalytically active self-splicing intron. We employed the algorithm of Tang et al.^12^ to scramble the ELP sequence to mitigate repetitiveness, but even still, RNAfold^31,32^ predicted that in these initial designs (called “LT1-ELP” for loopable translator 1 for elastin-like polypeptide), not all secondary structural elements on the intron properly form. Specifically, the critical P1, P3, P4, and P5 segments basepair with complementary segments in the ELP coding sequence (Fig. 1b, properly paired intron sequence in cyan, improperly paired sequences in yellow (intron) and blue (coding sequence)). Indeed, we found that when this RNA is transcribed in vitro, it does not circularize efficiently (∼6%), as based on a gel-shift assay and its inability to generate an RNase R-resistant product (Fig. 1c; N.B.: RNase R is an exonuclease). Initially, we reasoned that the “offending” ELP sequences that paired with the intron machinery could be excised, resulting in the “LT1-ELPmed” and “LT1-ELPshort” ELP constructs, but these also failed to circularize efficiently in vitro (Fig. 1c), presumably because of the large number of potential pairing partners in this GC-rich sequence (Supplementary Fig. 2a-b). When we annealed the RNA at 55 °C with 10 mM MgCl_2_, which promotes refolding of the intron, the circularization reaction proceeds and results in intensified lower bands (Fig. 1c). In follow-up experiments, we verified circularization by performing RT-PCR reactions that would only result in amplification on a circularized template (due to “outward-facing” primers, Supplementary Fig. 3a-c). Our observations suggest that misfolding of the intron’s catalytic core in the context of a GC-rich coding sequence is the source of the inefficiency.

Following previous work by Wesselhoeft et al.^33^, we next “insulated” the intron machinery (Fig. 1d, gray) from the coding sequence through two strong “locker” sequences (Fig. 1d, red). Whilst the external locker is removed along with the intron during circularization, the internal locker would create an undesired insertion in the coding sequence in the context of looped translation where ribosomes transit iteratively along the circular mRNA (Fig. 1d). To address this issue, we developed a Monte Carlo algorithm that identifies synonymous codon mutations to generate ELP-coding segments that would strongly basepair, thereby melding the internal locker sequence into the ELP-coding sequence (Fig. 1d–e; Supplementary Fig. 4). According to RNAfold, these locker sequences allow all the secondary structural elements of the intron’s catalytic machinery to properly form in the context of the whole 1600-nt primary transcript (Fig. 1f–g). We also scanned the two internal locker sequences through the remainder of the ELP coding sequence and made synonymous mutations to ensure they cannot also strongly basepair with other segments of the construct (Supplementary Fig. 4). By gel-shift assay, the resulting construct (called “LT2-ELP”) elevates the efficiency of circularization from 6% to 40% (Fig. 1i, lanes 1–2). We moreover confirmed that LT2-ELP circularizes mRNA by sequencing following RT-PCR with outward-facing primers (Supplementary Fig. 3d).

RNA helicases and RNA chaperones have been shown to facilitate folding of group I introns. Although their role has not, to the best of our knowledge, been systematically explored as an engineering component in synthetic circular RNA systems, we hypothesized that they could also promote spliced intron folding and therefore improve RNA circularization efficiency. Annealing RNA at 55 °C would not be compatible with an *in vivo* system since it is a lethal temperature for *E. coli*, but we reasoned an RNA helicase like CsdA^34^ or an RNA folding catalyst like *N. crassa* CYT-18 could replicate its effect (Fig. 1h)^35,36^. We expressed and purified these enzymes from *E. coli* (Supplementary Fig. 5), and found that at low (3 mM) MgCl_2_ concentrations, CsdA and CYT-18 did improve circularization efficiency of the LT2-ELP to 65% and 72% respectively (from 58%; Fig. 1i, lanes 6,8,10). Strikingly, the efficiency of circularization could be raised to high levels (>90%) when CsdA and CYT-18 were combined with higher (but physiologically relevant) concentrations of Mg^2+^. We conclude that the lockers and sequence optimization used in LT2, combined with RNA folding chaperones and divalent cations, can facilitate efficient circularization of repetitive, GC-rich mRNAs.

### Programming Ribosomes to Synthesize Ultra-Long Fibrous Proteins

In many bacteria, ribosomal translation commences at an internal Shine-Dalgarno (SD) ribosome-binding sequence (RBS)^37,38^, suggesting that they might initiate translation efficiently on circular mRNA and synthesize repetitive protein sequences by iteratively translating the same mRNA multiple times if there is no in-frame stop codon (Fig. 2a). *E. coli* and *B. subtilis* were transformed with plasmids encoding LT2-ELP, grown to log phase, lysed with detergent, and the whole extract was resolved by Tris-Acetate (3–8%) SDS-PAGE and visualized by anti-FLAG Western Blot. *E. coli* robustly translates FLAG epitope-containing proteins with molecular weights (MW) well above 460 kDa, which would require >15 transits around the circular mRNA (Fig. 2b). *B. subtilis* also expresses high-MW proteins above background, albeit with lower yield. The specific banding pattern in the Western blot suggests that ribosomes cease translating ELP in response to specific molecular events, rather than because of nonspecific termination or hydrolysis. It is now generally accepted that ribosome collisions serve as a major stress marker in both bacteria^39^ and eukaryotes^40^, and are recognized by ribosome quality control factors to promote mRNA decay or split stalled ribosomes, functions that are played by SmrB and HrpA in *E. coli*^41^. We reasoned that ribosome traffic and collisions could be principal factors that limit the yield of protein production from circular mRNA, given that in this environment ribosomes can initiate translation but cannot easily terminate (Fig. 2c).

**Fig. 2.**
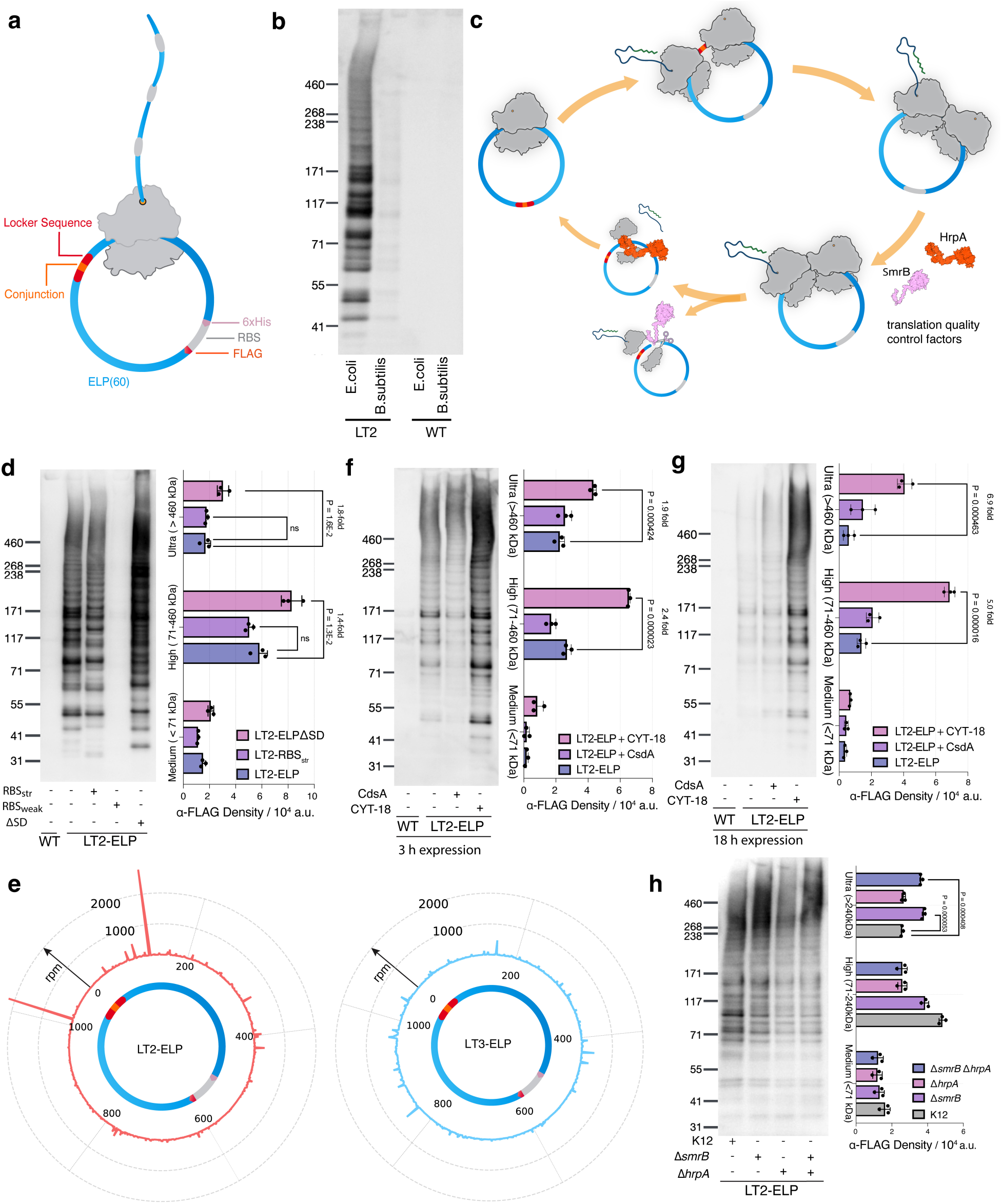
Looped translation. **a**, Illustration of looped translation on the circular RNA LT2. **b**, Anti-FLAG western blot assessing looped translation in *E. coli* NiCo21(DE3) and *B. subtilis* K07 harboring plasmids encoding LT2-ELP. WT, untransformed cells. **c**, Illustration of ribosome collisions and two rescue pathways in *E. coli*: HrpA pathway and SmrB pathway. **d**, The effect of varying RBSs and coding sequences on looped translation in *E. coli* as measured by anti-FLAG western blot of total cell extract. Protein expression was carried out for 4 h. Biological triplicates (*n* = 3) were transformed with the plasmid constructs indicated. For each replicate, identical levels of total protein were loaded. Densitometry was used for quantification, and P-values according to two-tailed unpaired t-tests are indicated. Representative western blot of one replicate shown. **e**, Ribosome footprint density as measured by Ribo-Seq when LT2-ELP and LT3-ELP circular mRNAs are expressed in *E. coli* NiCo21(DE3). **f**, The effect of RNA circularization-promoting factors on looped translation in *E. coli* as measured by anti-FLAG western blot of total cell extracts. Biological triplicates (n = 3) were transformed with the plasmid constructs indicated. For each replicate, identical levels of total protein were loaded. Densitometry was used for quantification, and P-values according to two-tailed unpaired t-tests are indicated. Representative western blot of one replicate shown. **g**, Analogous to panel f but with longer (18 h) induction times. **h**, The effect of ribosome collision rescue factor knock-out on looped translation in *E. coli* measured by anti-FLAG western blot of total cell extracts. Densitometry was used for quantification, and P-values according to two-tailed unpaired t-tests are indicated.

To test these theories, we modulated the ribosome binding site, which would control the rate at which new ribosomes are recruited onto circular mRNA to initiate translation. As LT2-ELP’s initial design possessed a conventionally strong SD sequence (de novo DNA RBS score 4270) we tuned it to a weaker RBS (with score 704) or a stronger RBS (LT2-RBS_str_, score 67735) (Supplementary Fig. 6). A weak RBS sharply reduced expression level, but unlike what would be expected with linear RNAs, strengthening the RBS did not improve expression levels either (Fig. 2d). This finding can be rationalized by noting that “over-recruitment” of ribosomes on circular mRNA could result in congestion and increased collision. One feature that has been suggested to play a role in slowing down the rate of ribosomal elongation is internal SD-like sequences^42^, due to their capacity to interact with 16S rRNA. Based on this concept, we redesigned the coding sequence of ELP by re-scrambling the codons subject to the constraint that internal SD-like sequences (e.g., all 6-nt sliding windows along the sequence with a calculated affinity to the anti-Shine-Dalgarno in excess of 8 kcal mol^-1^) are abolished (Supplementary Fig. 7a–b). The resulting construct, LT2-ELP(ΔSD), resulted in significantly higher levels of ELP expression, especially at the high (1.4-fold) and ultra-high MW ranges (1.8-fold, Fig. 2d). The improvement for ultra-high MW ranges became even more apparent (3.2-fold) when the expression persisted for 18 h (Supplementary Fig. 8a). To assess whether this boost could be attributed to mitigating ribosome traffic, we performed RiboSeq (i.e., ribosome profiling^43,44^) experiments on these two *E. coli* strains. We found that indeed LT2-ELP circular mRNA has two positions with very high levels of ribosome footprint density, but these features vanish when circular mRNA has internal SDs removed (Fig. 2e). Notably, these two sites with high ribosome density are 60 and 50 nt away from annotated SD-like sequences (Supplementary Fig. 7c–d). We refer to the version of the loopable translator lacking internal SD-like sites as LT3-ELP.

Because the availability of amino-acid-tRNA affects ribosome translation rates, we supplemented the expression culture with different concentrations of amino acids G, V, A, and P. However, we did not observe any apparent improvements in translation (Supplementary Fig. 8d). Altering the growth temperature also did not result in significant differences (Supplementary Fig. 8b). We also tested expression in the presence of rifampicin, which suspends the endogenous RNA polymerase activity but allows T7 polymerase to continue to synthesize the pre-circular ELP-encoding mRNA. We hypothesized that rifampicin would increase the circular RNA-to-ribosome ratio and thereby increase the chance of one ribosome per circular RNA. At a low concentration of rifampicin, we observed a modest increase in the loopable translation (Supplementary Fig. 8c). However, at higher concentrations, rifampicin was toxic to the cells. Finally, to further test the notion that ribosome collisions provide a mechanistic explanation for limited efficiency of looped translation, we transformed constructs into *E. coli* strains deficient in ribosome quality-control factors (Δ*smrB* and Δ*hrpA*, Fig. 2h). Deletion of SmrB modestly improves efficiency, whereas HrpA does not, suggesting that SmrB (which has endonuclease activity) may be responsible for mediating decay of circular mRNA with ribosome collisions.

Based on our *in vitro* observations establishing that mRNA circularization efficiency can be increased with helicases and folding catalysts, we next sought to test whether these improvements would be transferable to *E. coli*. We transformed cells with plasmids encoding either CsdA or CYT-18 in addition to LT2-ELP. CYT-18 markedly improves the expression of high-MW ELP (Fig. 2f), an effect that is amplified even further if cells are allowed to express protein for longer time intervals (Fig. 2g). After an 18 h induction, cells with CYT-18 were able to generate 5-fold more ELP in the high-MW range and 7-fold more ELP in the ultra-high-MW range than cells without CYT-18 (Fig. 2g). On the other hand, co-expressing CsdA did not result in significant improvements (Fig. 2f–g), for which we suggest two possible reasons: i) *E. coli* already endogenously express CsdA to a moderate level^34^, so cells may already be able to benefit from RNA helicase activity to help revert misfolded introns; ii) Overexpression of CsdA could reduce secondary structures that protect circular RNAs from endogenous endonucleases. Finally, we verified that LT2-ELP and LT3-ELP mRNAs are still circularized *in vivo*, by isolating RNA from cells via the TRIzol method and performing RT-PCR with outward-facing primers (Supplementary Fig. 9). These experiments showed that both LT2-ELP and LT3-ELP circularize at qualitatively similar levels, supporting the view that the substantive improvements in protein expression associated with LT3-ELP are due to translational efficiency. We conclude that through rational design, efficient translation of ultra-high MW ELPs can be achieved through looped translation in *E. coli*.

### Characterization of Ultra-High Molecular Weight ELP

To better estimate the molecular weight of the ELPs synthesized by LT3, we isolated protein from *E. coli* by lysing with detergent, supplementing with 8 M urea to improve solubility, then performing Ni-NTA affinity chromatography (that will bind the His-tag incorporated into the ELP), and dialysis. In shaker-flask batch cultures grown at 37°C, typical ELP yields were between 23–35 mg/L (Supplementary Fig. 10). Previous literature focusing on the largest known protein, Titin N2A, which has a nominal molecular weight of 3.8 MDa, demonstrated that SDS-agarose gels are amenable for resolving the largest proteins^45^. We first performed SDS-agarose electrophoresis on rat muscle homogenates and performed Coomassie staining to confirm that our conditions could resolve and visualize the highest MW proteins in rat muscle tissue (Supplementary Fig. 11), including nebulin (∼860 kDa), titin T2 (∼2.0 MDa) and titin N2A (∼3.8 MDa). Next, we electrophoresed purified ELP alongside homogenized rat muscle, transferred to a PVDF membrane, and immunoblotted with both anti-His and anti-Titin (Fig. 3a). The resulting protein (Fig. 3a, lane 1) resolved on SDS-agarose as a smear below Titin (lane 6), suggesting the typical molecular weight of the ELP polymer is on the order of 1–2 MDa. Given the heterogeneity of molecular weights for ELP, to see what the upper limit of the MW distribution is, we enriched for the highest MW products by ultracentrifuging the purified ELP at 55,000 rpm and electrophoresing the pellets. This fraction once again showed a smear, but shifted to higher MWs, with a portion of the density (16.2%) possessing an electrophoretic mobility even lower than that of titin N2A (Fig. 3a, lane 2). Hence, some of the elastin-like polymer produced by *E. coli* equipped with LT3 may be among the largest single-chain polypeptides synthesized by ribosomal translation.

**Fig. 3.**
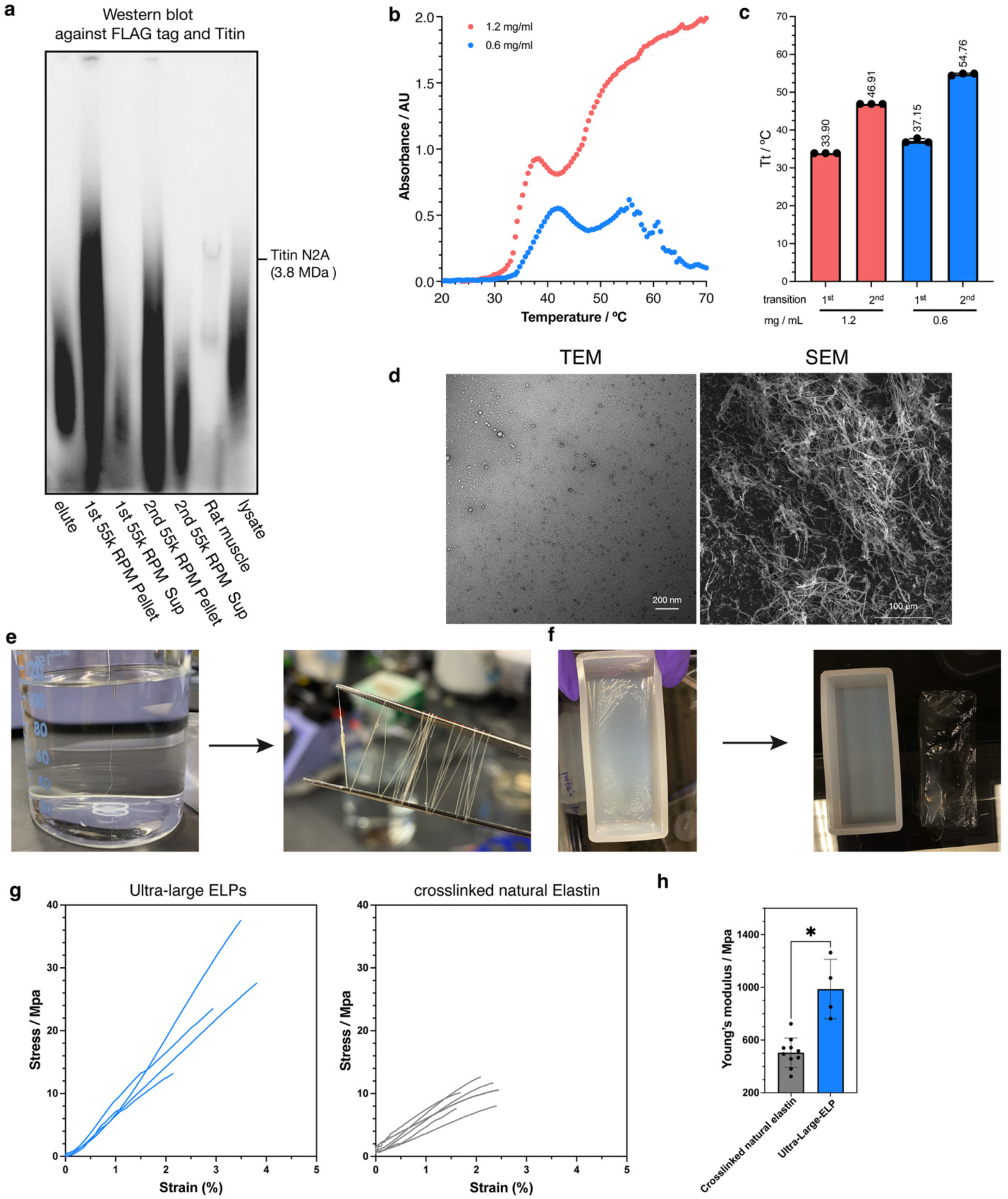
Characterization of ultra-high molecular weight ELPs produced through looped translation. **a**, Molecular weight comparison between titin and ELPs using protein agarose gel followed by western blot with anti-Titin and anti-FLAG antibodies. Purified ELPs were ultra-centrifuged to enrich ultra-high molecular weight species. **b**, Lower critical solution temperature measurement on the purified ELPs by solution turbidity. Technical triplicates (*n* = 3) for each concentration were measured. Representative curve shown. **c**, Bar chart showing measurements of transition temperature (*T*_t_). The slope of the turbidity curves were calculated, and *T*_t_ is defined as the temperature with the maximum instantaneous slope. **d**, Representative images from scanning electron microscopy (SEM) and transmission electron microscopy (TEM) on the purified ELPs. Scale bars provided. **e**, Protein wet-spinning process to fabricate an ELP fiber. **f**, Protein casting in a mold to fabricate an ELP film. **g**, Mechanical measurements of ELPs and crosslinked natural elastin. Protein films of ELPs synthesized through looped translation were measured in a tensile testing mode through dynamic mechanical analysis. The crosslinked fibers of natural elastin from bovine neck ligament were measured through tensile testing. **h**, The Young’s modulus of ultra-high molecular weight ELP and natural elastin from independent measurements (*n* > 4). *, P < 0.05 from unpaired two-tailed Student t-test.

One property of elastin-like polypeptides is that they exhibit lower critical solution temperatures (LCSTs)^46^, meaning that they are soluble at low temperature but insoluble at higher temperature. According to the Chilkoti model, for ELPs with a common sequence, the transition temperature becomes lower for ELPs with longer chain lengths (reciprocally) and higher concentrations (logarithmically). The polymer (GAGVPGVGVP)_60_ (which is the unit sequence of the product of LT3) has a critical temperature of 55.2°C at 0.6 mg/ml^12^. As expected, the product of LT3 has a significantly lower transition temperature of 37.1°C (Fig. 3b-c) at the same concentration, implying a higher average molecular weight. Moreover, the transition temperature decreases to 33.9°C at 1.2 mg/ml concentration (Fig. 3b-c), as qualitatively predicted by the Chilkoti model^46^. We note, however, that the presence of a second transition is a distinct behavior not found for (GAGVPGVGVP)_60_ and may be a consequence of the sample’s polydispersity (cf. Fig. 2d, 3a).

To further characterize the purified ELPs translated through looped translation, we carried out transmission electron microscopy on polymer solutions and scanning electron microscopy on lyophilized solids. The ELP forms micelles with an average diameter of 16 nm in solution, consistent with previous studies (Fig. 3d)^47^. However, they form nanofibers upon dissolution when becoming solids (Fig. 3d).

An important property of protein biomaterials is their ability to be fabricated into fibers or films. We sought to use a wet-spinning process to extrude ELP dope in HFIP solvent into a coagulation bath, successfully generating fibers (Fig. 3e). Additionally, the ELP dope could be readily cast onto molds and formed transparent robust films upon air-drying (Fig. 3f). When the same process was carried out on bovine serum albumin (BSA), proteins were unable to form such films (Supplementary Fig. 12). We used dynamic mechanical analysis to measure the mechanical properties of ultra-high MW ELP films (Fig. 3g-h), which showed a Young’s modulus at 987.2±225.6 MPa. In contrast, natural elastin proteins purified from bovine neck ligament were neither able to form films through casting nor could they be converted into stable fibers through extrusion into a coagulation bath. To attempt to create a control to compare ultra-high MW ELP to, we applied 0.1% glutaraldehyde to the wet-spinning process of natural elastin. Interestingly, even with the crosslinking of 0.1% glutaraldehyde, natural elastin fibers have a Young’s modulus of only 505.0±111.6 MPa on average (Fig. 3g-h). Thus, ultra-long ELPs can provide access to otherwise inaccessible material properties.

### LT3 Generalizes to Other Fibrous Protein Sequences

Much of the original design behind LT2 was prompted by the constraints imposed by the fact that elastin’s glycine-, proline-, and alanine-rich sequence (encoded by GGN, CCN, and GCN codons) has very high %GC, complicating mRNA circularization and iterative ribosomal translation. This constraint is not unique to elastin: Other important natural fibrous proteins, like collagen ((GP[A/P])_n_)^48^, keratin (CCXPX, CCX[S/T][S/T] repeats)^49^, and silk^50^, also make extensive use of the same amino acids encoded by GC-heavy nucleotide sequences. Hence, we next examined whether the strategies that we implemented in LT2 to improve circularization and in LT3 to improve translation were robust to promote these two processes on other similarly challenging but functionally relevant sequences.

We therefore applied the rational criteria that went into designing LT3-ELP (e.g., codon scrambling, internal locker sequence generation and melding, SD-like sequence removal) to design LT3’s for five further fibrous proteins: DHF58^51^, collagen-like protein ((GAPGTPGPQGLPGSP)_n_)^12^, (AEAEAKAK)_n_^52^, (RADA)_n_^53^, C3HR^7^ as well as a repetitive ankyrin domain^54^. Based on computational predictions, all the designs could properly fold the secondary structures of the intron in the context of the full mRNA (Supplementary Fig. 13), and four of them had internal SD-like sequences abolished (Supplementary Fig. 14). The designed sequences were cloned into a plasmid with T7 promoters and transcribed *in vitro* with T7 RNA polymerase. We found that they were all able to circularize with at least moderate efficiency (Fig. 4a). EAK peptide circularized the most efficiently at 59.5%, comparable to what we found for ELP before supplementing with CsdA or CYT-18 (40%). As previously (cf. Fig. 1), to establish the identity of the lower bands as the circularized RNA, we found that in all cases the lower band alone survives RNase R treatment, and that its intensity is increased when subjected to annealing at 55 °C with 10 mM MgCl_2_. The LT3 constructs were transformed into *E. coli*, all expressing respective repetitive protein materials with a His tag. Lysis of the cells followed by Western blot showed that all LT3 constructs produced high MW proteins, with one exception being the poly-Ankyrin motif protein (Fig. 4b). Under longer exposure times, Western blotting showed that concatemer forms of poly-Ankyrin motif protein were generated, though with relatively low yield, which could be due to its low RNA circularization efficiency. Given that these systems used identical promoters, RBS, and plasmid copy number, it is apparent that some repetitive proteins (particularly (EAK)_n_ and (RADA)_n_) express even more readily than ELP. We conclude that the design elements incorporated into LT3 provide a general solution to circularize a diverse set of repetitive GC-rich mRNA sequences and translate high-MW polypeptide materials from them.

**Fig. 4.**
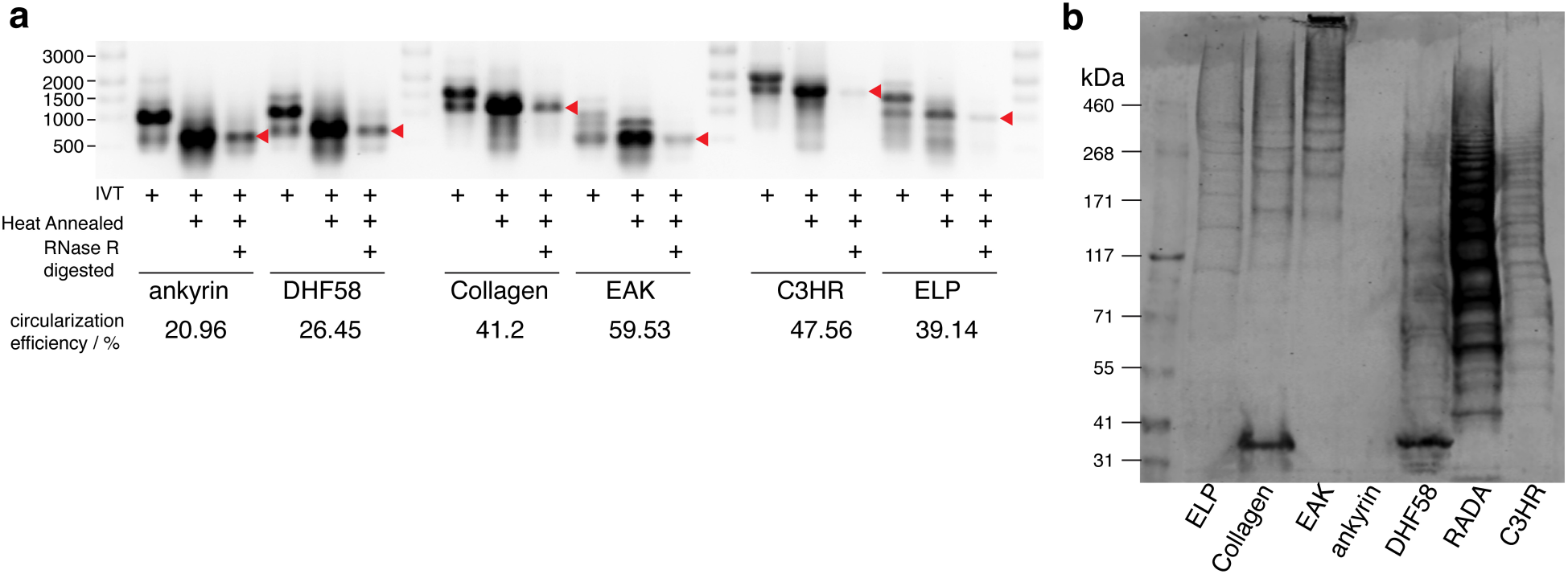
The generalization of looped translation for different repetitive fibrous proteins. **a**, Electrophoretic mobility shift assays and RNase R assays on *in vitro* transcribed/circularized RNA constructs corresponding to diverse fibrous proteins, with/without re-annealing. Red arrows denote the circularized product. **b**, Anti-FLAG western blot for looped translation from *E. coli* NiCo21(DE3) charged with LT3 systems programmed to translate looped versions of seven different fibrous proteins indicated, all labeled with the FLAG epitope.

### Coupling Looped Translation to Secretion through the Translocon

High-MW proteins that are trapped in a bacterium’s cytoplasm are unlikely to be available for processing and downstream use; we hence considered potential ways to couple ribosomal translation to co-translational export through the SecYEG translocon^55^, also known as the general export pathway (Fig. 5a-1). In particular, because SecYEG can interface with the ribosome directly and nascent chains transit through its protein-conducting pore synchronously with their translation, we reasoned that this translocon could be uniquely well suited for secreting ultra-high MW proteins whose translation “never” ends.

**Fig. 5.**
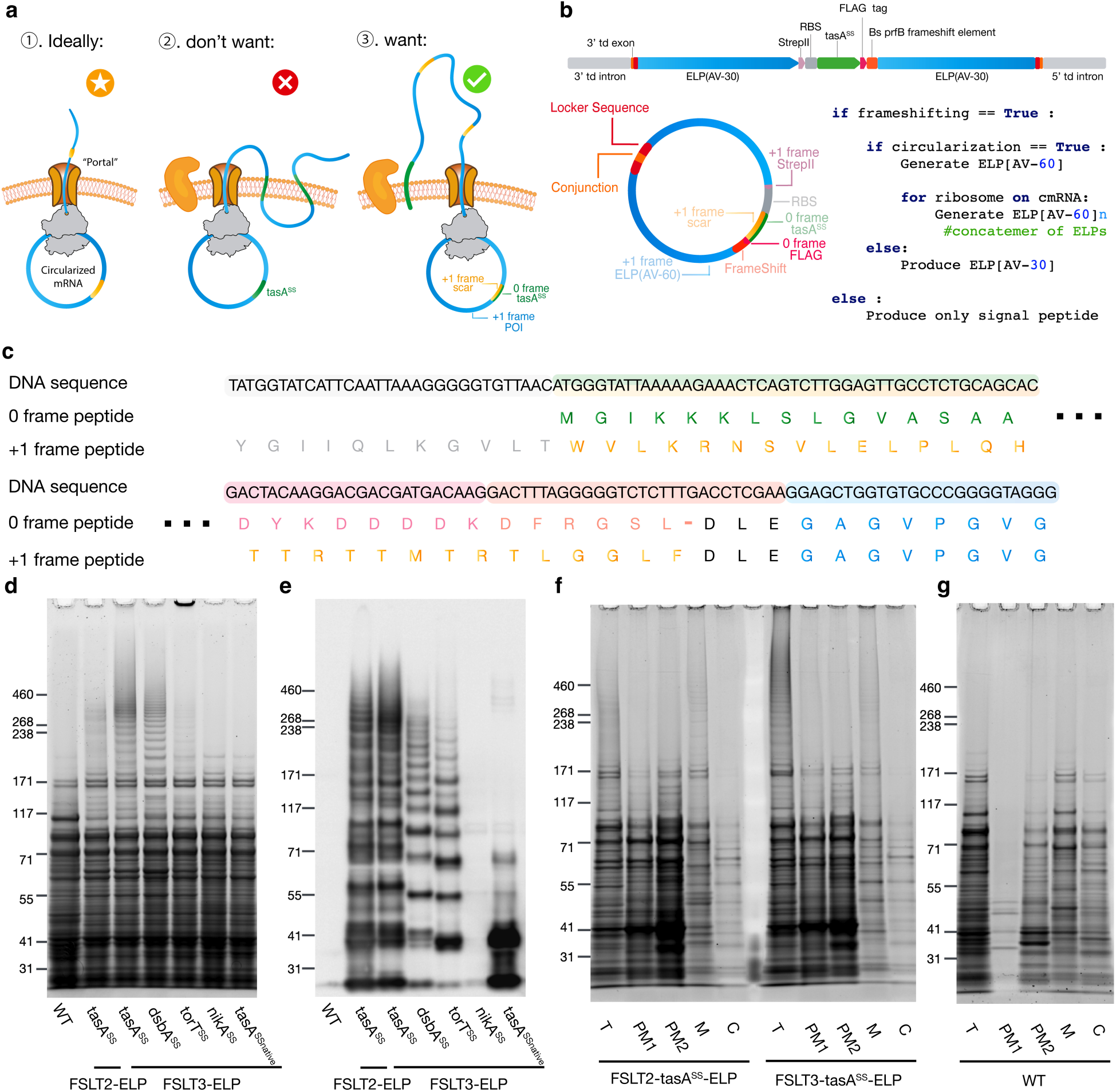
Looped secretor for production and secretion of ultra-high molecular weight polypeptides. **a**, Illustration of loopable secretion and the importance of translating the signal sequence once. **b**, Design of looped secretor and translation logic schemes and an if-tree showing the possible outcomes of the synthetic mRNA system depending on whether the RNA is circularized and/or frameshifting occurs at the programmed ribosomal frameshift (PRF) element. **c**, Sequence of the RF2 +1 frameshifting element in the context of circular RNA. **d**-**e**, Total protein gel (SyproRuby stained, **d**) and western blot (anti-FLAG, **e**) of *E. coli* cells expressing looped secretor using different signal peptides. **f**, Subcellular localization of polypeptides translocated across the inner membrane as assessed by SyproRuby staining of the indicated fractions. The fractions are total protein (T), 1^st^ periplasmic fraction (PM1), 2^nd^ periplasmic fraction (PM2), membrane fraction (M), and cytosol fraction (C). **g,** Similar to f but for untransformed cells.

Canonically, the SecYEG translocon is engaged by a hydrophobic signal sequence (SS) that is recognized by either the signal recognition particle (SRP) or SecA^56,57^. While we^58^ and others^59^ have demonstrated that certain signal sequences can be requisitioned to direct the secretion of fibrous proteins, a fundamental challenge emerges in the context of trying to secrete looped proteins through this mechanism: It would be undesirable to “loop over” the signal sequence, since these hydrophobic helical segments typically partition into membranes which would likely result in a polytopic membrane protein (Fig. 5a-2) instead of continuous secretion; multiple signal peptidase cleavage sites could lead to fragmentation of ultra-high MW polypeptides. To design a looped translator that only translates the signal sequence once (Fig. 5a-3), we turned to programmed ribosomal frameshifts (PRFs, Fig. 5b)^60^. Specifically, we reasoned that if we inserted a PRF in between the signal sequence and the ELP-encoding sequence (and placed ELP in a +1 frame), then the ribosome upon reprising onto the original mRNA segment encoding the signal sequence in the +1 frame would instead translate a linker sequence that is neither helical nor particularly hydrophobic, so as to allow continuous extrusion through SecYEG rather than partitioning into the plasma membrane (Fig. 5b–c). To create this construct, which we call FSLT2-tasA^SS^-ELP (or FSLT3-tasA^SS^-ELP, with SD sequences removed from ELP), three design elements were incorporated: (i) We prepended the ELP coding sequence with the SS from *B. subtilis* tasA and the slippery sequence from *B. subtilis* prfB (protein release factor 2); (ii) We installed on a separate promoter copy of SipW, a dedicated signal peptidase from *B. subtilis* that is specific to the tasA^SS^; and (iii) We edited the codons of tasA^SS^ to ensure that in the +1 frame it does not encode any stop codons and encodes a polypeptide that is disordered (according to AlphaFold2 prediction) with low hydrophobicity (Supplementary Fig. 15). In this construct, failure to frameshift results in a short 5.7 kDa product because the codon immediately following the PRF in the 0 frame is a stop codon; failure to circularize results in a 19.2 kDa (or 16.6 kDa if signal sequence is removed) product which corresponds to half of the ELP codons on the circular mRNA (because half of the ELP is upstream of the start codon, lower mass corresponds to with cleavage of the signal peptide); and failure to loop results in a 33.4 kDa product (or 30.8 kDa, if signal sequence is removed).

To test FSLT3-ELP, we cloned a small series of constructs in which the SS was varied (Fig. 5d-e), transformed them into *E. coli*, and analyzed the total protein content by SDS-PAGE, visualizing with SyproRuby (a non-specific stain, Fig. 5d) and anti-FLAG immunoblot (Fig. 5e). Untransformed cells (lane 1) did not produce any protein larger than 171 kDa, or any material that was responsive to FLAG antibody. On the other hand, two signal sequences, that from *B. subtilis tasA* and from *E. coli dsbA*, resulted in robust levels of high-MW protein (11.9% and 7.3% of total protein content for tasA^SS^ and dsbA^SS^, respectively, by densitometry (Supplementary Fig. 16a-b)) that can be confidently assigned to ELP based on the similarity of the band structure between SyproRuby stain and immunoblot (Fig. 5d-e, lanes 3-4). FSLT3-tasA^SS^-ELP appears to outperform FSLT2-tasA^SS^-ELP, and moreover, if we use the original codons of tasA^SS^ (in which stop codons would be present in the +1 frame), we obtain the major expected product at 41 kDa, corresponding to the result if circularization succeeds, but looping fails due to termination upon the first reprisal through the SS. A low-MW band below 31 kDa, found on all constructs in this Tris-acetate SDS gel, likely represents the two products resulting from either failed frameshifting or circularization. In separate Tris-glycine gels, which resolve lower MW proteins (Supplementary Fig. 17), we confirmed that a common failure point is circularization.

Next, to test that FSLT3-tasA^SS^-ELP is capable of both synthesizing and secreting translocating high-MW fibrous proteins, we reperformed these protein expressions but subjected the lysates to a subcellular fractionation protocol^61^, which separates *E. coli*’s cytoplasm (C), from the inner plasma membrane (M), and periplasm (PM1 and PM2). The subcellular fractions were resolved by SDS-PAGE and visualized with SyproRuby along an unfractionated, total (T), sample (Fig. 5f-g). In untransformed cells (Fig. 5g), there is no protein detected with MW greater than 171 kDa, and very little protein detected in the PM1 fraction. Cells equipped with FSLT3-tasA^SS^-ELP, on the other hand, synthesize high-MW proteins that are found in the periplasmic and membrane fractions, and are absent from the cytoplasmic fraction (Fig. 5f). We therefore conclude that FSLT3-tasA^SS^-ELP coordinates loopable translation with co-translational secretion to direct *E. coli* to translocate high-MW repetitive fibrous proteins out of the cytoplasm.

### *Bacillus subtilis* exports protein with a frameshifting loopable translator

In our earlier experiments (cf. Fig. 2b), we found that LT2-ELP was capable of expressing protein in *B. subtilis* as well as *E. coli*, albeit with lower yields. We were motivated then to see whether the improvements developed in *E. coli* to evolve LT2-ELP into FSLT3-ELP would also translate into a system that could export ultrahigh molecular weight protein directly into the growth media, given that in Gram-positive bacteria, the translocon is localized in the plasma membrane and there is no further membrane barrier external to it. Initially, we screened two signal sequences that were the most successful in *E. coli*: tasA^SS^, which produced the highest density of bands at high molecular weight, and torT^SS^, which produced less protein between 170-460 kDa, but intriguingly produced a band with very low electrophoretic mobility (cf. Fig. 5d). We also interrogated two types of growth media and two induction times (Fig. 6a). tasA^SS^ resulted in very little protein passing into the secreted fraction (e.g., supernatant, S), with most staying within the cellular (C) fraction. On the other hand, FSLT3-torT^SS^-ELP resulted in significant levels of proteins detected in the supernatant, especially after extended induction periods (Fig. 6a). The ultra-high molecular weight polymers which we observed in *E. coli* were not observed in *B. subtilis*, suggesting that this bacterium’s translation and secretion apparatus may impose additional constraints on loopable secreted translation. The strongest features near 55 and 117 kDa suggest the looped secretor could pass the circular RNA 1 to 4 rounds and export the protein into the extracellular environment.

**Fig. 6.**
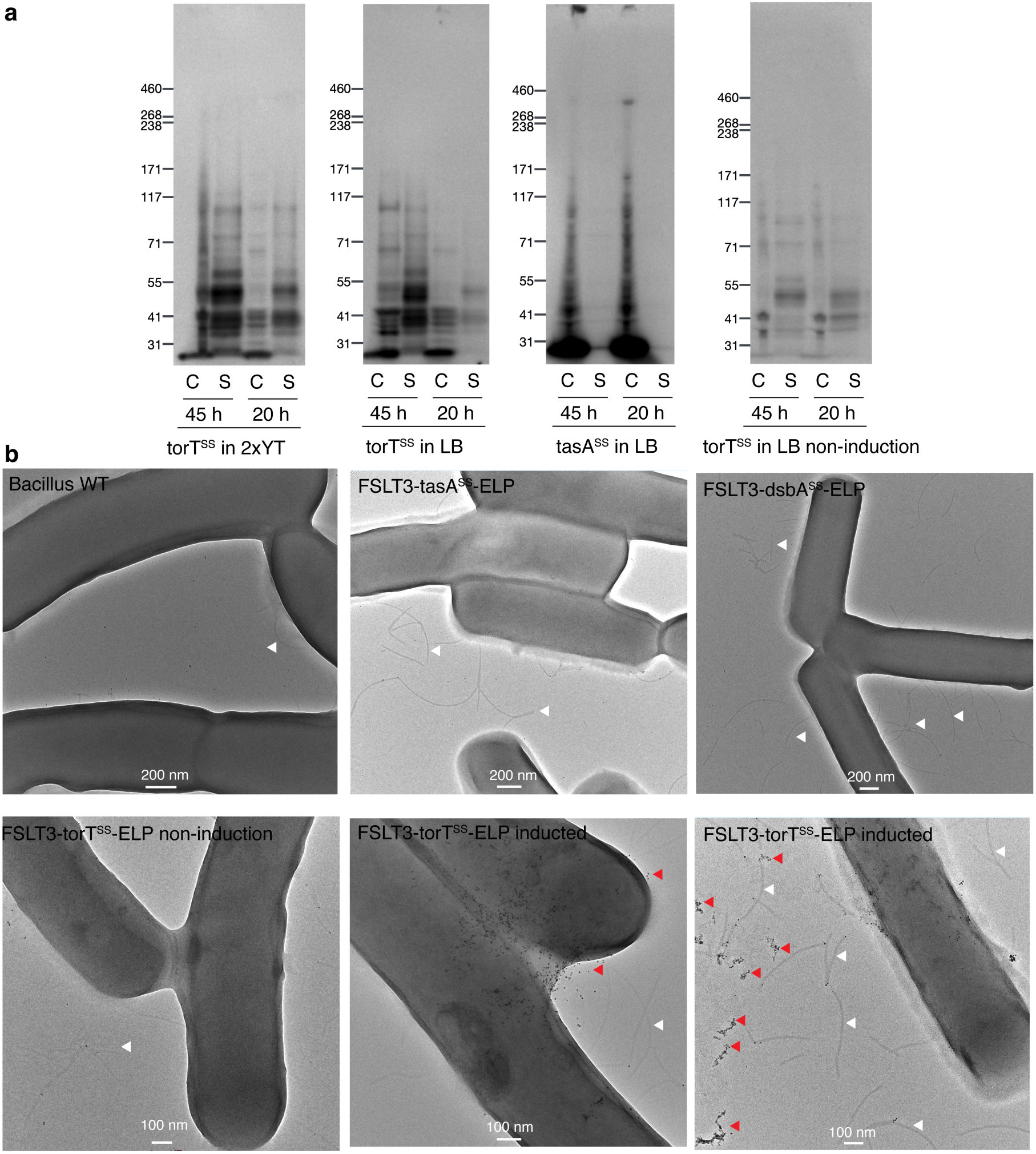
Looped secretor in the context of a Gram-positive bacterium, *Bacillus subtilis*. **a**, Anti-FLAG Western blot results of *Bacillus subtilis* expressing a looped secretor with different signal peptides. The expressions were carried out in different media (2xTY, LB) with two different time periods (20 h, 45 h). The cellular (C) and secreted (S) fractions were separated by centrifugation at 4000 g for 20 mins. Only the top 10% of the supernatants were collected for western blot. Cell pellets were washed with PBS to remove medium. **b**, Nano-gold labeled transmission electron microscope images of *Bacillus subtilis* expressing loopable secretor with different signal peptides. Large rods are *Bacillus subtilis* cells. White arrows indicate the flagella from the bacteria. Ni-NTA nano-gold particles (5 nm, red arrow) exhibit dark black particles that colocalize with secreted ELP polymer fibers.

We also performed transmission electron microscopy studies on *B. subtilis* cells harboring various FSLT3-ELP machineries with distinct signal sequences (Fig. 6b). Encouragingly, we found that when *Bacillus* equipped with FLST3-torT^SS^-ELP are induced, the cells are visualized with fibrillar emanations that also stain positively for His-tag based on immunogold labeling using gold nanoparticles coated with the nickel-NTA epitope. Fibrillar structures detected in transmission electron micrographs of all *B. subtilis* strains are typical shapes of flagella^62^, which did not stain with the immunogold nanoparticles as well. Hence, we conclude that FSLT-torT^SS^-ELP equips *B. subtilis* to express and simultaneously secrete ELP polymers with high molecular weight into the growth media. However, additional investigation and optimization will be necessary to achieve the ultra-high molecular weights observed in Gram-negative bacteria.

## Discussion

Proteins with ultra-high molecular weights have unique mechanical properties, but because of the size of the genes necessary to encode them and the specialized machinery to process them, they are typically associated with metazoans that possess contractile muscle fibers. On the other hand, microorganisms tend to be practical systems for recombinant protein expression and production because of how readily they can be cultured and genetically manipulated. There are several applications that can benefit from microbial production of fibrous ultra-high molecular weight proteins, including cultured meats, sustainable materials (like biological equivalents to concrete and binders), and tissue engineering. A number of challenges have, however, limited our ability to program microbes to produce ultra-high molecular weight proteins. Synthesis of repetitive DNA sequences is oftentimes very challenging^63^ and can also be unstable with respect to recombination in microbial hosts. Moreover, the presence of large fibrous polymers in the cytoplasm can be toxic to many organisms.

In this report, we have sought to overcome these limitations by developing a robust platform to circularize protein-coding RNA sequences to enable loopable translation and demonstrate that this biotechnology can be coupled to programmed ribosomal frameshifting to enable secretion of proteins with higher molecular weight. The idea of using the permuted intron method to circularize mRNA is not new, however, the challenges we encountered with generalizing it to repetitive fibrous protein sequences and the absence of many publications using the method following its initial proof-of-concept are suggestive that significant challenges prevented the permuted intron method from being practically useful for a range of important applications. Through a series of mechanistically guided engineering strategies, we established design principles that substantially improve RNA circularization by 15-fold and looped translation of fibrous protein from undetectable to 30 mg/L titer. Our final design incorporates a range of synergistic features including the inclusion of: i) synonymous locking architecture that both isolates the ribozyme structure from adjacent repetitive coding sequence and preserves protein identity; ii) addition of RNA helicase CsdA and RNA chaperone CYT-18; iii) removal of SD-like sequences that slow down ribosomes. These numerous considerations become significant for circularizing sequences with high GC-content and repetitive character, which are generally the case for fibrous protein modules. We find that the optimized loopable translator design we report here, LT3, is robust and can efficiently circularize RNAs and express ultra-high molecular weight protein for six out of seven sequences that we tested. The addition of a SecYEG signal sequence that is only translated once in a unique frame from the rest of the encoded sequence moreover allows robust secretion of ultra-high molecular weight proteins in *E. coli*. In *E. coli,* secretion from the translocon results in localization into the periplasm. In the periplasm, ultrahigh molecular weight proteins may be less toxic to the host cell and can also be retrieved by permeabilization of the outer membrane. Moreover, there are many secreted proteins in industry that are produced in *E. coli,* including human growth hormone, antibody fragments (Fabs), and therapeutic enzymes like L-Asparaginase^64^. We sought to replicate our same success in *B. subtilis*, where secretion from the translocon results in localization to the extracellular environment. While this system is still capable of circularizing RNA, and performing loopable translation, it does not secrete proteins with ultrahigh molecular weights, and hence requires further optimization. It is known that translation initiation is quite different in *B. subtilis* owing to the absence of a ribosomal protein bS1, and that it uses a very distinct pathway to resolve collided ribosomes^65^. These considerations may change the requirements for efficient loopable translation in *B. subtilis*.

In conclusion, we present tools and strategies to program microbes to produce and secrete ultrahigh molecular weight polymers. Under favourable conditions, proteins produced with these tools can exceed the molecular weight of titin N2A from mammalian muscle, making them amongst the largest single-chain polypeptides produced by a ribosome. Given the robustness of LT3 with respect to sequence, the methods demonstrated here are well stationed to enable the synthesis and evolution of programmable protein materials from microbial hosts.

## Methods

### General materials and strains

Bacterial strains and plasmids used in this study are listed in Table S1. All enzymes for molecular biology used in this study were purchased from New England Biolabs, USA (NEB) and Thermo Fisher Scientific, USA. All primers were obtained from Integrated DNA Technologies, Inc. (IDT). Standard Sanger sequencing was performed by Genewiz. Whole plasmid sequencing was performed by Plasmidsaurus. *E. coli* was grown in lysogeny broth (LB) medium, 2xYT medium, or terrific broth (TB) medium (as indicated) supplemented with ampicillin (100 μg/mL) or kanamycin (100 μg/mL) when needed. *B. subtilis* strains were grown in LB medium or 2xYT medium (as indicated) supplemented with 1 μg/mL erythromycin and 25 μg/ml lincomycin when needed.

### Cloning of plasmids

#### Cloning of construct LT1-ELP

Gene fragment with the repetitive sequence of ELP as well as the split intron sequences were designed and ordered through the Genscript service. The fragment was amplified using primers Pr1 and Pr2 and inserted into the plasmid, pLIKE-sipW-tasAss(30)-silk(10Ala)1, which was derived from our previous work, through the Gibson Assembly strategy. The backbone was amplified using primers Pr3 and Pr4. The PCR products were supplemented with 0.5 µL DpnI for 30 minutes and subsequently purified by the Zymo DNA clean & concentrator kit according to the manufacturer’s protocols. The size of the purified DNA was checked by gel electrophoresis, and the DNA concentration was measured by a NanoDrop One microvolume UV-Vis spectrophotometer (Thermo Scientific). The Gibson assembly was carried out using HiFi DNA assembly master mix (NEB) according to the manufacturer’s protocol. The mixture was transformed into NEB 10-beta chemically competent *E.coli* cells and plated on LB agar plates supplemented with 100 µg/mL ampicillin. Colonies formed following overnight incubation at 37 °C were picked and transferred into 5 mL LB liquid culture with 100 µg/mL ampicillin. Mini-preps were carried out to extract the plasmids, which were verified by Sanger sequencing service by the Genewiz company. The validated plasmid was denoted as LT1-ELP.

#### Cloning of LT1-ELP derivatives LT1-ELP_med_ and LT1-ELP_short_

The plasmid construct LT1-ELP was used as a template for PCR using primer pairs: Pr5 and Pr6 for LT1-ELP_med_; Pr5 and Pr7 for LT1-ELP_short_. The PCR products were processed as described above but then transformed directly into 10-beta chemical competent *E.coli* cells in a Quick-change cloning manner. The plasmids were validated by Sanger sequencing.

#### Cloning of LT2-ELP and its derivatives – RBSstr, RBSweak and ΔSD

Gene fragment of LT2-ELP ordered through Genscript was amplified using primer pair: Pr8 and Pr9. The fragment was inserted into the pLIKE template through Gibson Assembly strategy. The template was amplified using primer pair: Pr10 and Pr11. The Gibson Assembly procedure was carried out as described above. The validated plasmid, denoted as LT2-ELP, was then used as a template for cloning of RBSstr and RBSweak using primer pairs: Pr12 and Pr13 for RBSstr, Pr14 and Pr15 for RBSweak. ELP(ΔSD) sequences were first designed and pre-optimized using the customized codon scrambling tool to reduce repetitiveness and avoid SD-like sequences at the nucleotide level. Its gene fragment was then ordered through the Genewiz gene synthesis service. The sequence was amplified using primer pair: Pr16 and Pr17, and inserted into the plasmid through Gibson Assembly, in which the template of PCR was LT2-ELP using primer pair: Pr18 and Pr19. The resulting plasmid was denoted as LT3-ELP.

#### Cloning of repetitive fibrous sequences of DHF58, collagen-like protein, EAK RADA, C3HR and ankyrin

Gene fragments were ordered through the Genewiz gene synthesis service. The repetitive protein sequences were pre-optimized using the codon scrambling tool, developed by Tang et al, to reduce the repetitiveness at the nucleotide level. In addition, the codons of DHF58, collagen-like protein, EAK RADA, C3HR, and ankyrin were scrambled to avoid SD-like sequences by customizing the codon scrambling tool. The gene fragments were inserted into the plasmid same as ELP(ΔSD) described above.

#### Transformation of plasmids into *Bacillus* spp

Electrocompetent *Bacillus subtilis* cells were prepared following an optimized electroporation protocol. Briefly, strain K07 was grown overnight in 5 mL LB medium at 30 °C for 14 h to reach late logarithmic phase (OD_600_ = 1.0–1.5). Overnight cultures were diluted 1:10 into 50 mL of fresh 2×YT medium supplemented with 0.5 M sorbitol and grown at 37 °C with shaking until OD_600_ reached 0.8–1.0 (approximately 3.5 h). Cultures were centrifuged at 4,000 × g for 5 min at 4 °C. Cell pellets were resuspended in ice-cold washing solution (0.5 M sorbitol, 0.5 M mannitol, and 10% glycerol). The washing procedure was repeated a total of four times. After the final wash, cells were resuspended in 2.5 mL washing solution. For electroporation, 200 µL of electrocompetent cells were mixed with 200 ng of plasmid. Electroporation was performed at 2.5 kV with a pulse length of 5.0 ms. Immediately after pulsing, 900 µL of pre-warmed recovery medium (2×YT supplemented with 0.5 M sorbitol and 0.38 M mannitol) was added, and cells were recovered at 37 °C for 3 h with shaking. Cells were plated onto LB agar containing appropriate antibiotics. Plates were incubated overnight at 37 °C.

#### Protein expression and purification

Colonies carrying the looped translation constructs were grown overnight in LB medium at 37 °C. The overnight cultures were then inoculated into 2 liters of fresh TB medium at an initial OD_600_ of 0.05. When OD_600_ reached 0.4, the looped translation was induced by adding IPTG (Isopropyl β-D-1-thiogalactopyranoside) to a final concentration of 1 mM. Cultures were shifted to 30 °C and incubated overnight.

Cells were harvested by centrifugation at 4000 g for 30 min at 4 °C and resuspended in 200 mL lysis buffer (50 mM Tris pH 8.0, 150 mM NaCl, 1% Triton X-100, 0.1% SDS, 0.5 mg/ml lysozyme, 0.1 mg/ml DNase I, 1xProtease Inhibitor, 2 mM MgCl_2_). After incubation at 37 °C, urea solids were added to a final concentration of 8 M to clarify the lysate. The lysates were incubated with 10 mL of Ni-NTA resin at 4 °C for 1 hour in the presence of 5 mM imidazole. Afterward, the resin-lysate mixtures were passed through a column to collect the resin. Beads were washed with 25 mL of washing buffer (50 mM Tris pH 8.0, 300 mM NaCl, 8 M urea, 10 mM imidazole) until A_280_ absorbance approached 0.01(8-9 washes). The polypeptides were eluted from the beads with 10 mL of elution buffer (50 mM Tris pH 8.0, 300 mM NaCl, 8 M urea, 300 mM imidazole) for 5 rounds. The solution was dialyzed stepwise against buffer A (25 mM Tris pH 8.0, 150 mM NaCl, 4 M urea), buffer B (25 mM Tris pH 8.0, 150 mM NaCl), and buffer C (100 mM ammonium bicarbonate). The final solution was lyophilized on a FreeZone Freeze Dryer (operated at-48 °C and 0.01 mBar, Labconco).

#### SDS-PAGE using Tris-acetate gels and western blot

Bacteria expressing the constructs under different conditions (as indicated in the main text) were centrifuged to produce a pellet. The pellets were then mixed with BugBuster master mix (Novagen) supplemented with cOmplete protease inhibitor (Roche). For Bacillus spp., additional lysozymes were added with a final concentration of 0.5 mg/ml. The lysates were then incubated at 37°C for 10 minutes until the lysates were cleared. To ensure the lysate was completely clarified and large aggregates were removed, SDS was added with a final concentration of 0.5%.

To ensure equal loading of protein for analysis, the total protein concentrations of the lysate were measured using Pierce BCA Protein Assay (Thermo Scientific) following standard protocols. Then, the total protein concentrations were diluted and normalized. 26 µl of normalized total proteins were mixed with 4 µl DTT (1M) and 10 µl NuPAGE LDS sample buffer (Thermo Scientific). The mixture was incubated at 70 °C for 10 minutes, chilled on ice for 5 minutes, and loaded onto NuPAGE Tris-acetate mini protein gels (3 to 8%). The electrophoresis was conducted using Tris-acetate SDS running buffer (50 mM Tricine, 50 mM Tris Base, 0.1% SDS, pH 8.24) at 30 mA and 4 °C for 90 minutes.

After electrophoresis, electroblotting was performed using an iBlot 2 Gel Transfer Device (Thermo Fisher) according to the manufacturer’s protocol (3 min at 20 V; 4 min at 23 V; 3 min at 25 V). The PVDF membranes were incubated in 5% non-fat milk in TBST buffer (20 mM Tris, 150 mM NaCl, 0.1% Tween 20, pH 7.6) for 1 hour at room temperature. Next, the PVDF membranes were incubated with primary antibody (anti-His or anti-FLAG antibody as indicated, Invitrogen), which was diluted 1,000x in TBST buffer with 5% non-fat milk. After overnight incubation at 4 °C, the PVDF membranes were washed with TBST buffer 3 times (10 mins each time). The PVDF membranes were then incubated with the secondary antibody (anti-mouse-HRP antibody, Invitrogen), which was diluted 10,000x in TBST buffer with 5% non-fat milk for 1 hour at room temperature, followed by washing using the TBST buffer. For imaging, the chemiluminescence reagents (Signal West Femto Maximum Sensitive Substrate; Thermo Fisher Scientific) were mixed in a 1:1 ratio for 1 min with a total volume of 800 μl and then applied to the PVDF membranes. The images were acquired using a Chemi-Doc imager (BioRad).

To confirm equal loading, PVDF membranes post-imaging were washed with TBST buffer and ultrapure water each 3 times. PVDF membranes were then stained with Ponceau S Staining Solution (Thermo Fisher) for 15 minutes. PVDF membranes were washed with ultra-pure water to reveal the protein bands, which were imaged using a Chemi-Doc imager (BioRad).

#### Titin extraction, Protein Agarose Gels and Electroblotting

Titin protein extraction and protein agarose gel for the analysis of ultra-large proteins were conducted following the protocol by Chad M. Warren and Marion L. Greaser. Briefly, rat muscle tissue was mixed with sample buffer (8 M urea, 2 M thiourea, 3% SDS, 75 mM DTT, 0.03% bromophenol blue, and 0.05 M Tris, pH 6.8). The mixture was placed in a Dounce homogenizer to lyse the cells. The sample was then vortexed and heated for 10 minutes at 60 °C. The sample was then centrifuged, and the supernatant containing titin molecules was stored at-80 °C.

For 1% protein agarose gels, agarose powder was mixed with glycerol as well as buffers to make a final composition as: 30% glycerol, 50 mM Tris, 0.384 M glycine, 0.1% SDS. The gel was cast into empty Invitrogen Gel Cassettes (Thermo Fisher). After the gel was solidified at 4 °C for 1 hour. Samples were loaded onto the gel, and the gel electrophoresis was conducted using running buffer: 50 mM Tris, 0.384 M glycine, 0.1% SDS. The upper buffer chamber was supplemented with 10 mM 2-mercaptoethanol.

For electroblotting, protein agarose gels were placed in transfer buffer: 10 mM CAPS, pH 11, 0.1% SDS, 10 mM 2-mercaptoethanol. Following that, the gels were placed on PVDF membrane, and the blotting was conducted as described above. Anti-FLAG and anti-titin antibodies were used for the detection of both polyELPs and titin.

#### Ribo-seq analysis

Ribosome profiling was conducted through service by EIRNA bio. The data were processed and analyzed according to the procedure by Fuad Mohammad and Allen Buskirk^66^. The data were converted into circular plots using code described in the source code.

#### In vitro transcription

In vitro transcriptions were conducted using HiScribe® T7 High Yield RNA Synthesis Kit (E2040S) following standard protocols. DNA templates using plasmids or PCR products were mixed with necessary kit components and incubated at 37°C for 4 hours. To remove template DNA after the reaction, 70 μl nuclease-free water, 10 μl of 10X DNase I Buffer, and 2 μl of DNase I (RNase-free) were added, mixed, and incubated for 15 minutes at 37°C. Synthesized RNAs were purified through column purification (Zymo Research’s RNA Clean & Concentrator) following standard protocol.

#### Enhanced RNA circularization

For the heat annealing method of circularization, RNA was heated to 70 °C for 5 min and then immediately placed on ice for 3 min. Then, GTP was added to a final concentration of 2 mM along with a buffer including magnesium (50 mM Tris-HCl, 10 mM MgCl2, 1 mM DTT, pH 7.5). RNA was then heated to 55 °C for 15 min and then column purified.

#### RNase R digestions

20 μg of RNA was diluted in water (86 μL final volume) and then heated at 70 °C for 3 min and cooled on ice for 3 min. 20U RNase R and 10 μL of 10× RNase R buffer were added, and the reaction was incubated at 37 °C for 2 hours.

An additional 10U RNase R was added halfway through the reaction. RNase R-digested RNA was column purified.

#### RNA agarose gel

1 gram of TopVision agarose (Thermo Fisher) in 100 ml of 0.5x TBE buffer was boiled using a microwave to allow the agarose to dissolve. SYBR Safe Gel Stain was added after the solution cooled down but before solidifying. The 10.5 x 6cm gels were then cast using a mini gel mold.

RNAs were then mixed with 2X RNA Loading Dye (Thermo Scientific), incubated at 70 °C for 5 minutes, and chilled on ice for 3 minutes before being loaded into the mini gel. The electrophoresis was conducted using Accuris myGel Mini Electrophoresis System at 50 V.

#### Reverse-transcription PCRs

RT-PCR was conducted using SuperScript™ III reverse transcriptase (Invitrogen) following standard protocols.

#### Scanning electron microscopy

Lyophilized polypeptide solids were transferred onto a specimen stub with carbon tapes and imaged using the secondary electron mode on a Thermo Scientific Helios G4 UC Focused Ion Dual Beam Scanning Electron Microscope (Materials Characterization and Processing Core Facility, Johns Hopkins), operated at 5 kV accelerating voltage.

#### Transmission electron microscopy and immunogold labeling

Protein samples were applied onto carbon-coated copper grids (400 mesh, Electron Microscopy Sciences), which were glow-discharged by the Gloqube Plus Glow Discharge System operated at 1 mA plasma current and 5 kV. The grids were then washed with ultra-pure water 3 times and stained for 10 s in 50-μl droplets of UranyLess negative staining solution (Electron Microscopy Sciences) four times, with the carbon-coated face upside-down. The grids were then washed with ultra-pure water 3 times. The excess liquid on the grids was blotted off with filter papers (Fisherbrand course porosity, 9.0 cm diameter). The grids were allowed to air dry and were then examined using an FEI Talos F200C FEG TEM.

For immunogold labeling, 1 mL of culture following an overnight secretion expression was centrifuged at 8000 g, 4 °C for 10 min. The cell pellet was resuspended in PBS with 1% glutaraldehyde to fix the cells for 1 hour. After 3 washes using PBS, the cell pellets were incubated with PBS with 0.02 M glycine to quench glutaraldehyde. The cell pellets were then resuspended in PBS. After that, 10 μL resuspensions were applied onto carbon-coated copper grids with incubation for 10 minutes. The grids were then rinsed with ultrapure water three times, blocked with PBST-BSA (2%), then washed with PBST-imidazole (5 mM). The grids were placed face-down in a 100-μL droplet of PBST-imidazole with 10 nM 5 nm Ni-NTA-AuNP particles (Nanoprobes) to incubate for 90 min. The grids were then washed five times with PBST-imidazole, then twice with PBS. Post-fixing was conducted using a final concentration of 1% (v/v) glutaraldehyde in PBS for 15 minutes. Then remove the fixative by washing with PBS three times. The grids were finally rinsed in deionized water (2 × 1 mins), stained with Uranyless (5 × 1 min). Then rinse in deionized water (3 × 1 min). The grids were allowed to air dry and were then examined using an FEI Talos F200C FEG TEM.

#### Subcellular Fractionation of *E.coli* Cells

Isolation of subcellular fractions of *E.coli* cells was done based on the protocol by Kielkopf et, al. Briefly, 4 mL of bacterial culture was harvested by centrifugation at 5,000 × g for 15 min at 4 °C, and the supernatant was removed. To precipitate secreted proteins, ammonium sulfate was added to 3.2 mL of the collected LB supernatant to a final mass of 2.224 g, followed by incubation on ice for at least 20 min. The mixture was centrifuged at maximum speed for 20 min in a microcentrifuge, the supernatant was discarded, and the resulting pellet was resuspended in 500 µL of 1× PBS. In parallel, cell pellets were washed three times with 1 mL of ice-cold wash buffer (10 mM Tris, pH 8.0), centrifuging at 5,000 × g between washes to re-pack the cells.

Washed cell pellets were resuspended in 500 µL of cell lysis buffer containing 20% (w/v) sucrose and 30 mM Tris-Cl (pH 8.0) at room temperature, followed by addition of EDTA to a final concentration of 1 mM. The suspension was mixed gently for 10 min to chelate divalent cations and weaken the outer membrane. Cells were then pelleted by centrifugation at 10,000 × g for 10 min at 4 °C, and the supernatant was kept and denoted as PM1. The resulting pellets were thoroughly resuspended in 500 µL of ice-cold osmotic shock buffer (5 mM MgSO4) and incubated on ice for 10 min to induce osmotic shock. Following centrifugation at 10,000 × g for 10 min at 4 °C, the supernatant containing periplasmic proteins was collected and denoted as PM2. The remaining spheroplast pellet was resuspended in 500 µL of 0.1 M Tris-Cl (pH 8.0) and subjected to three cycles of freeze–thaw lysis by alternating between liquid nitrogen (dry ice) and incubation at 37 °C to gently lyse the spheroplast. Ultracentrifugation at 40000 g for 30 mins was done to separate the membrane fraction and cytosolic fraction. The membrane pellet was resuspended in 500 µL of buffer (20 mM Tris-HCl, pH 8.0, 5 mM EDTA, and 0.5% SDS).

## Data and Material Availability

All the data necessary to reproduce the study are found in the main text figures and supplemental figures. Annotated sequences used to generate the constructs described in the text are found in Table S3.

## Acknowledgements

We acknowledge helpful discussions with Yunsheng Sun and Sarah Woodson on RNA discussion and CsdA gift, Allen Buskirk for Riboseq analysis, and Beijun Shen for discussion on mechanical analysis. We are appreciable to Robert Louder at the Johns Hopkins Integrated Imaging Center and staff at the University of Delaware’s Advanced Materials Characterization Lab for technical support. S.D.F acknowledges support from NIGMS (R35-GM161721), an NSF CAREER grant (MCB-2045844), the Sloan Foundation, and the Dreyfus Foundation. M.D.S acknowledges funding support from NIGMS (R35-GM136354). A.M.B. was funded by an NSF Graduate Fellowship.

