## supplementary information for "Synthesis of Ultra-Large Fibrous Proteins from Bacteria via a Looped-Translation System"

[a] Current affiliation: Department of Molecular and Cellular Biology, University of California, Berkeley, CA 94720 USA

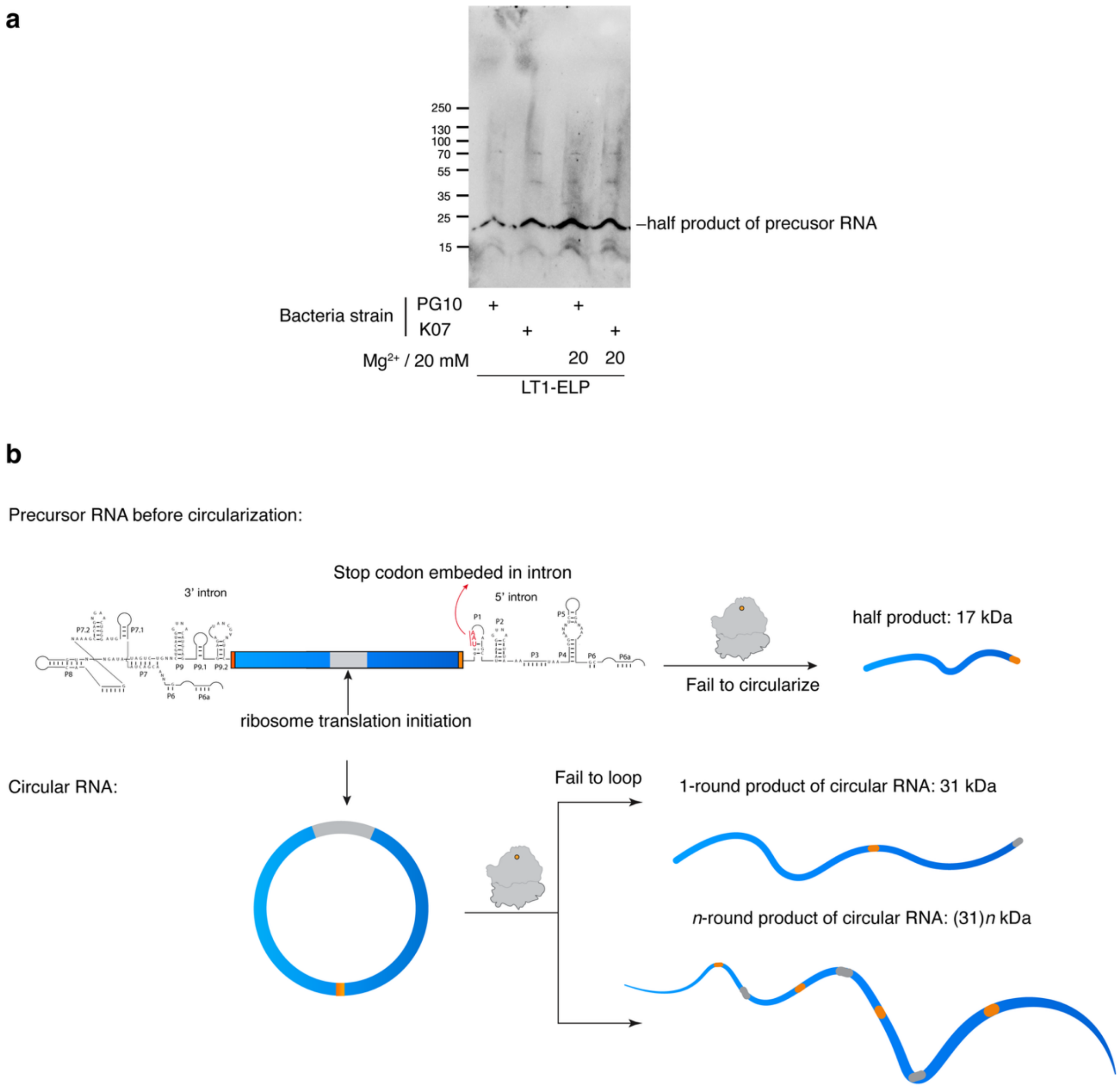

**Supplementary Figure 1**. **a**, Expression levels from the initial circular RNA design, LT1-ELP, as measured by 4- 20% Tris-glycine gel and anti-FLAG Western blot. K07 and PG10 are *B. subtilis* strains: K07 increases protein production with seven proteases knocked out; PG10 is a strain that underwent significant genome reduction[1,2]. **b**, Illustration of looped translation products. Precursor RNA will only produce a low molecular weight ELP at 17 kDa. Looped translation on circular RNA can result in concatemers of ELP proteins.

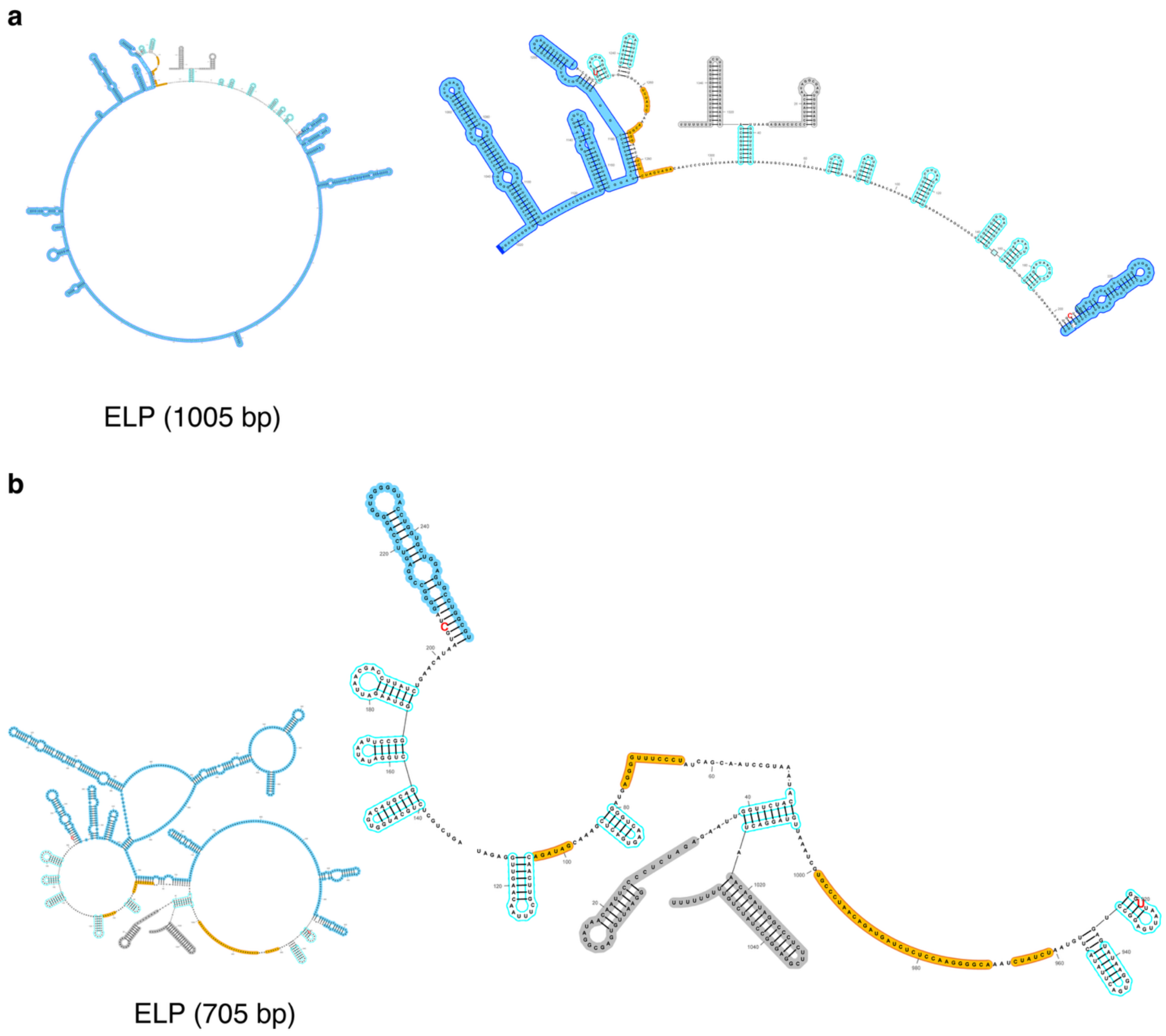

**Supplementary Figure 2**. RNAfold[3] minimum-free energy predicted secondary structures for LT1-ELP medium (**a**) and LT1-ELP short (**b**). The different lengths of ELP sequences were placed between the intron sequences. The overview of the predicted secondary structure is shown on the left side. The regions associated with the intron sequences are shown on the right side, zoomed in. Sequences of LacO and the terminator are colored in grey. Secondary structures of T4 intron sequences are colored cyan if pairing matches the desired secondary structure or in yellow if they are mispaired. Coding sequences are colored in blue.

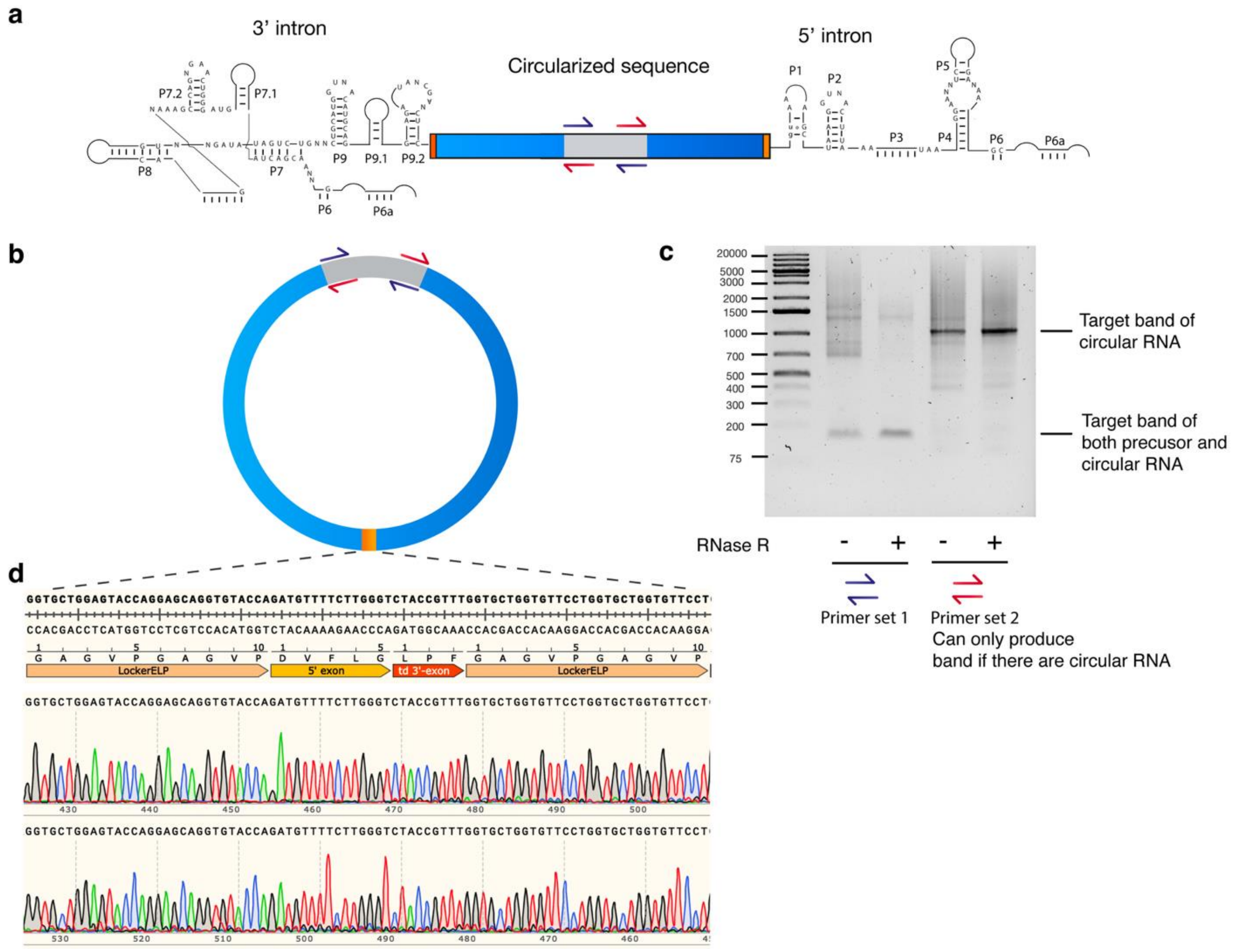

**Supplementary Figure 3**. Orthogonal characterization of circular RNA through reverse transcription-PCR (RT-PCR) and sequencing. **a-b**, Illustration of PCR primers to detect circularized mRNA in the context of the linear pre-circularized mRNA and the circularized mRNA product. **c**, Detection of circular mRNA synthesized through *in vitro* transcription, treated with or without RNase R. 1% agarose gel stained with SYBR Safe showing products from PCR with the primers indicated. **d**, Sanger sequencing runs showing continuous read through the junction site, proving correct circularization occurred between the expected nucleotides.

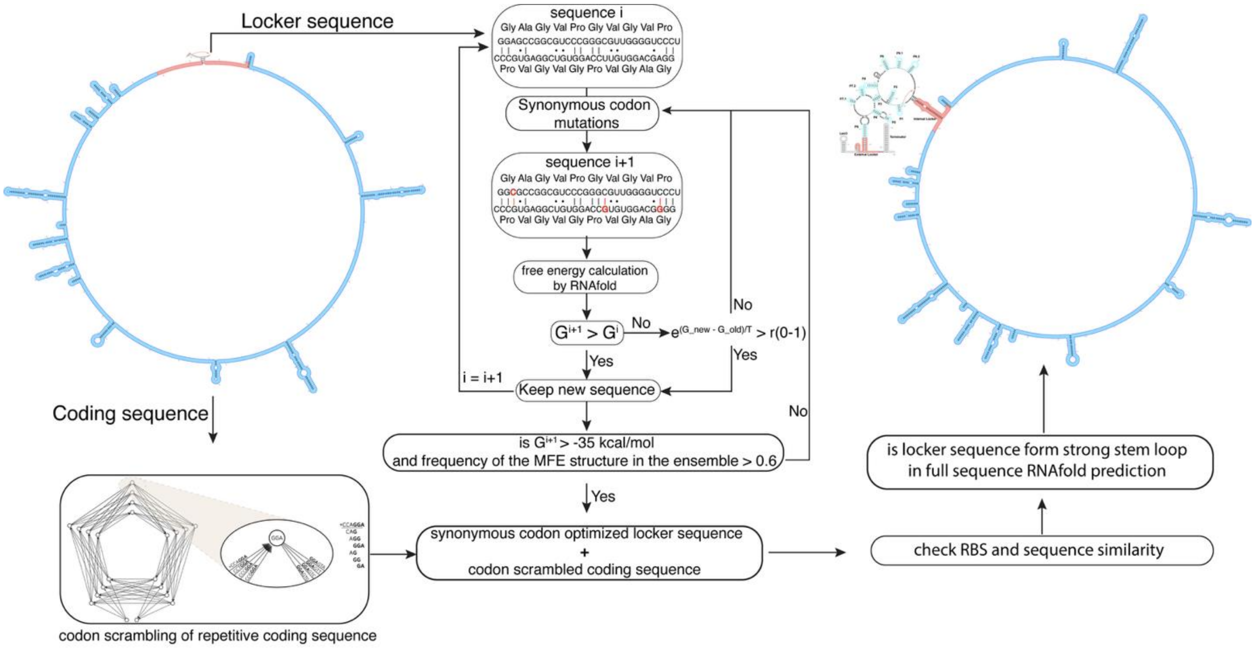

**Supplementary Figure 4**. Monte Carlo algorithm for the design of scarless internal locker sequences by melding them into the desired protein-coding sequence with synonymous codon engineering.

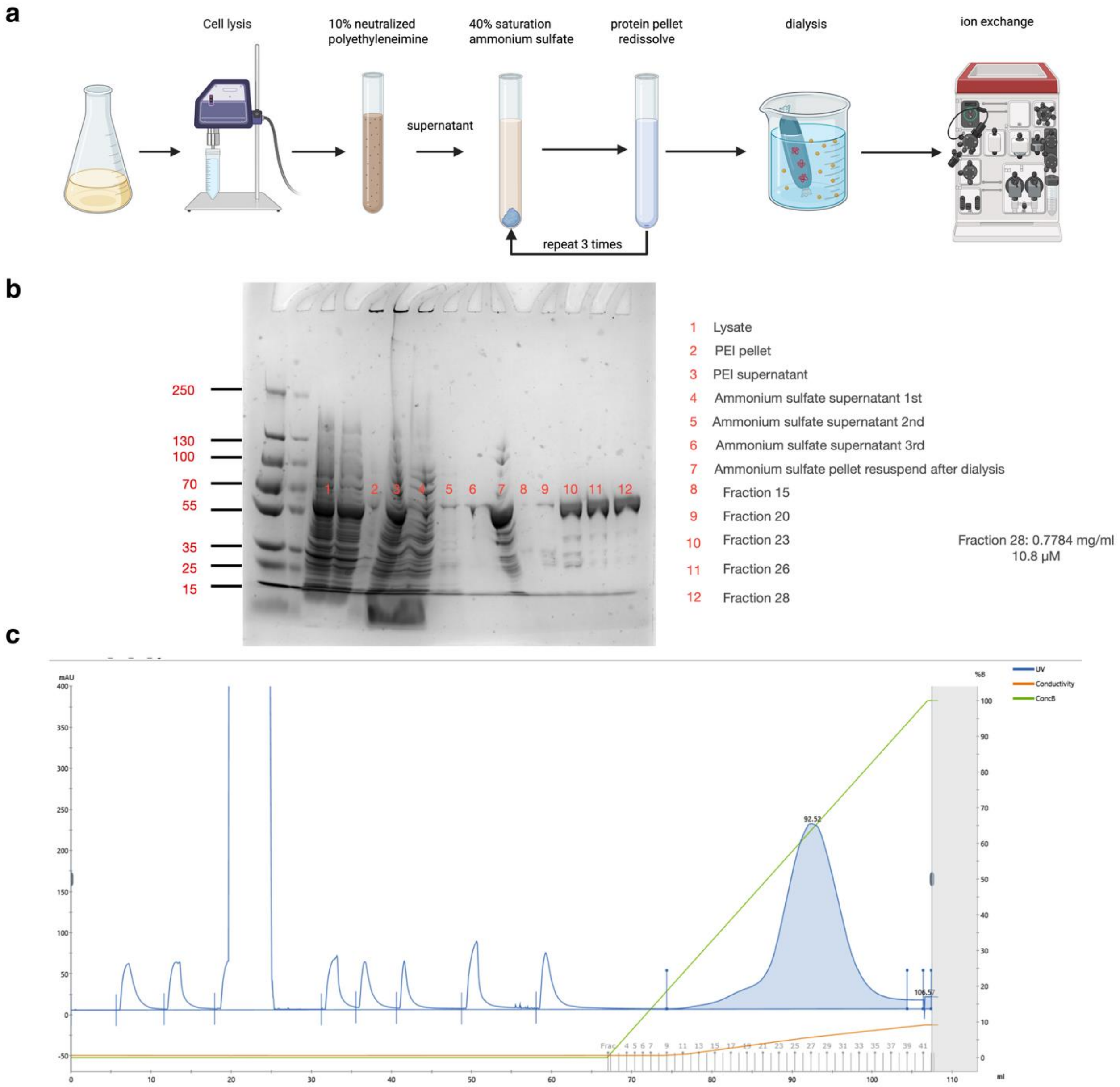

**Supplementary Figure 5**. Purification of CYT-18 by the PEI-precipitation procedure developed by R. Saldanha et al[4]. **a**, Overview of purification procedure. **b**, SDS-PAGE stained with Coomasie showing significant enrichment of CYT-18 following ammonium sulfate precipitation steps, and near purity achieved through ion exchange chromatography. **c**, Anion exchange chromatogram run to purify CYT-18.

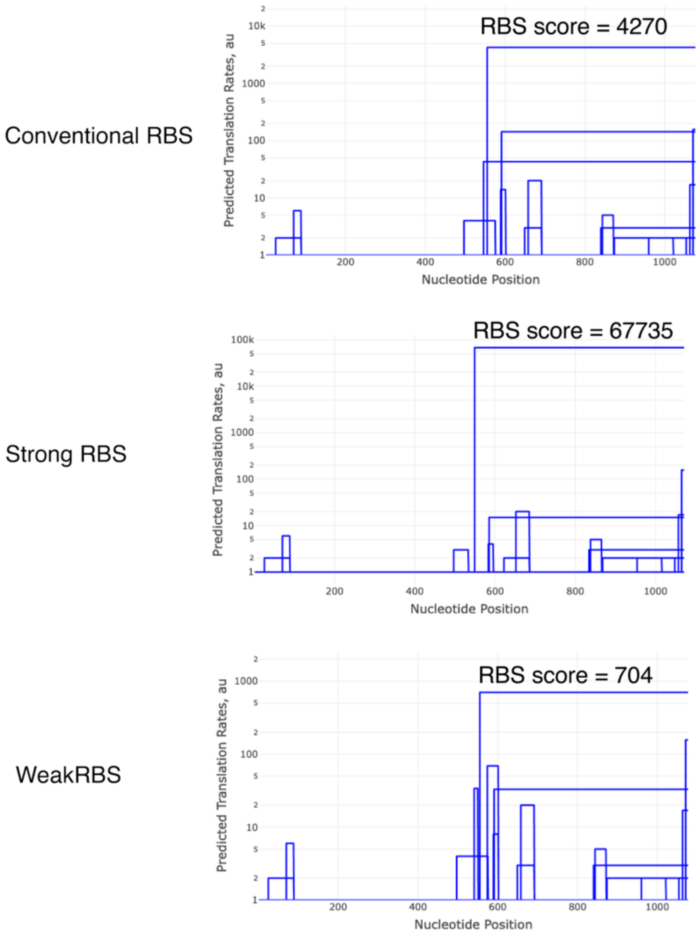

**Supplementary Figure 6**. Results from de novo DNA ribosome binding site (RBS) scoring calculator[5] for the original Shine-Dalgarno (SD) sequence used in LT2-ELP ("conventional") and two variants in which the SD sequence was either strengthened (strong RBS) or weakened (weak RBS).

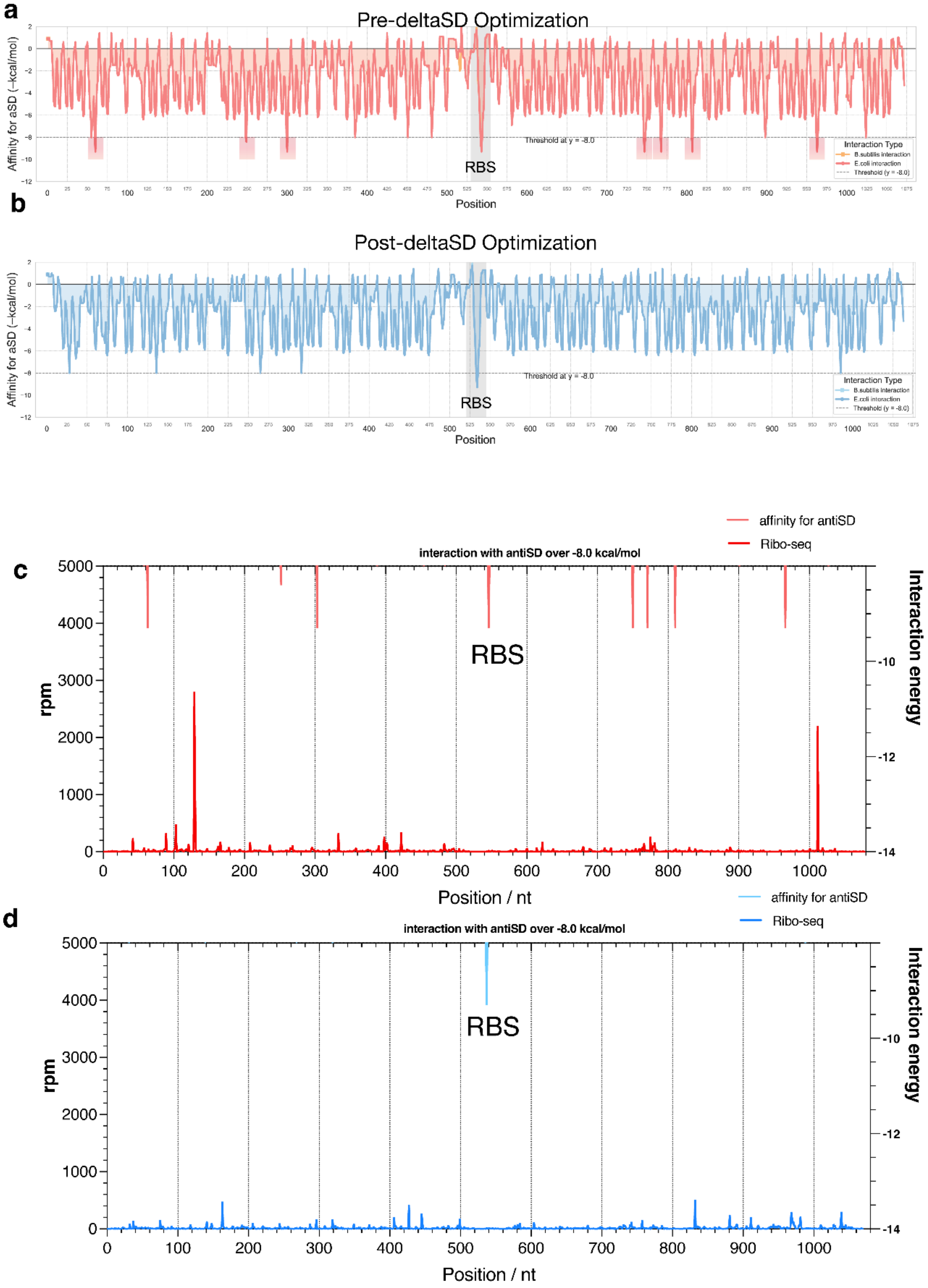

**Supplementary Figure 7**. Coding sequence interaction energy calculation probing for internal SD-like sequences. **a-b**, Interaction energy map of circular RNA sequence to the anti-Shine-Dalgarno sequence from both *E. coli* and *B. subtilis*. Note: the interaction data of *E. coli* and *B. subtilis* overlap with each other with *E. coli* data shown on the top. RBS sequences were highlighted in gray. Positions with interaction energy below -8 kcal/mol, which is characterized as a strong interaction by Li et al.[6], are highlighted. **c-d**, Combined view of interaction energy map and the Ribo-seq data.

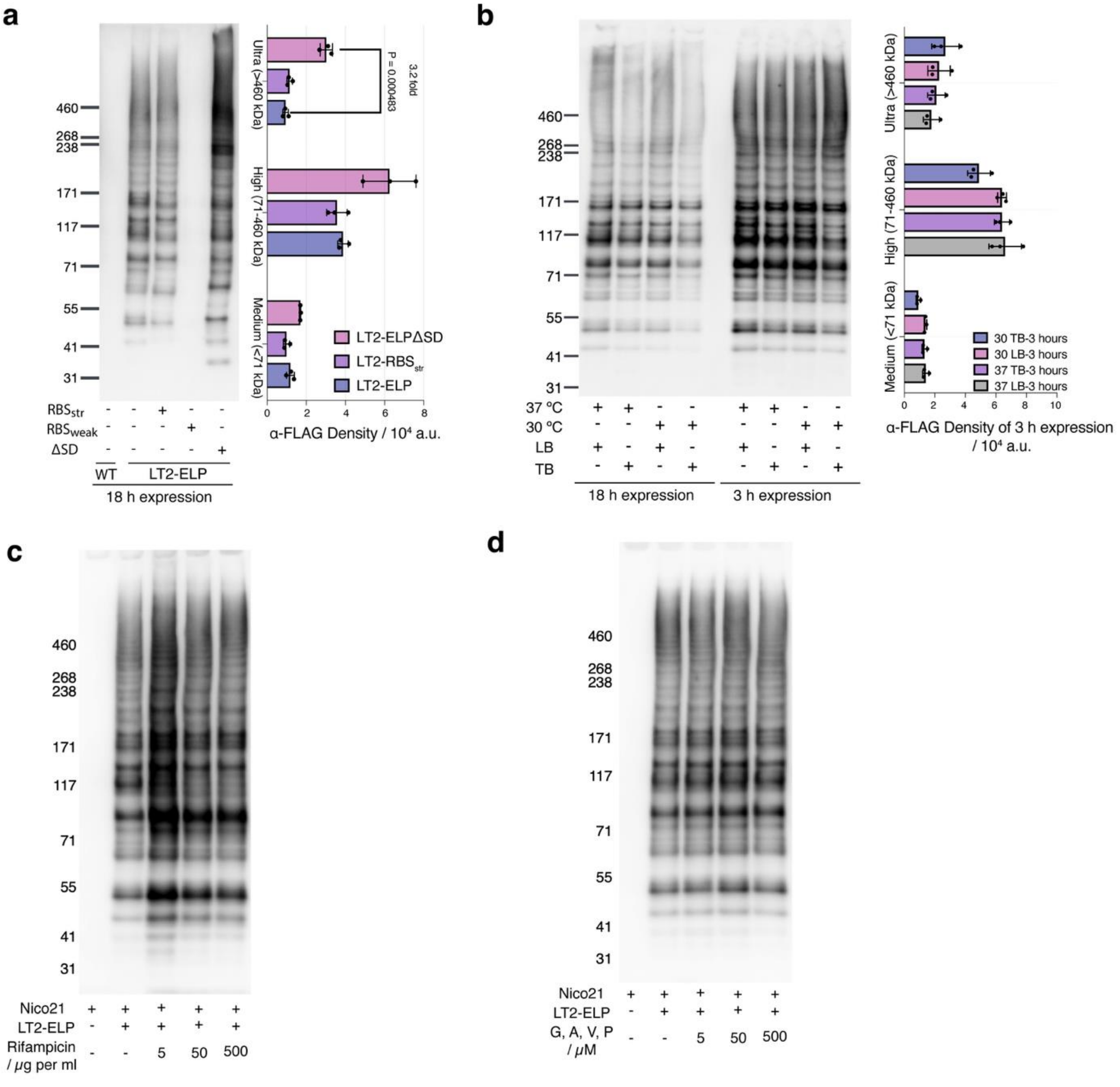

**Supplementary Figure 8**. Optimization of ELP expression conditions from circular mRNA. **a**, The effect of varying RBSs and coding sequences on looped translation in *E. coli* following 18 h expression as measured by anti-FLAG western blot of total cell extract. Biological triplicates (n = 3) were transformed with the plasmid constructs indicated. For each replicate, identical levels of total protein were loaded. Densitometry was used for quantification, and P-values according to two-tailed unpaired t-tests are indicated. Representative western blot of one replicate is shown. **b,** The effect of growth temperature and media type (LB or TB broth) on looped translation in *E. coli* following 3 h or 18 h expression as measured by anti-FLAG western blot of total cell extract. Biological triplicates (n = 3) were transformed with the plasmid constructs indicated. For each replicate, identical levels of total protein were loaded. Densitometry was used for quantification, and P-values according to two-tailed unpaired t-tests are indicated. Representative western blot of one replicate shown.

**c**, Effect of rifampicin on the looped ELP production as measured by anti-FLAG western blot. Total protein concentration was normalized, loaded on the same gel, and subsequently transferred onto the same PVDF membrane for antibody detection and chemiluminescent imaging. Nico21, a substrain of *E. coli* BL21(DE3).
**d**, Effect of amino acid supplementation on the looped ELP production as measured by anti-FLAG western blot. Total protein concentration was normalized, loaded on the same 3-8% Tris-Acetate gel and subsequently transferred onto the same PVDF membrane for antibody detection and chemiluminescent imaging. Nico21, a substrain of *E. coli* BL21(DE3).

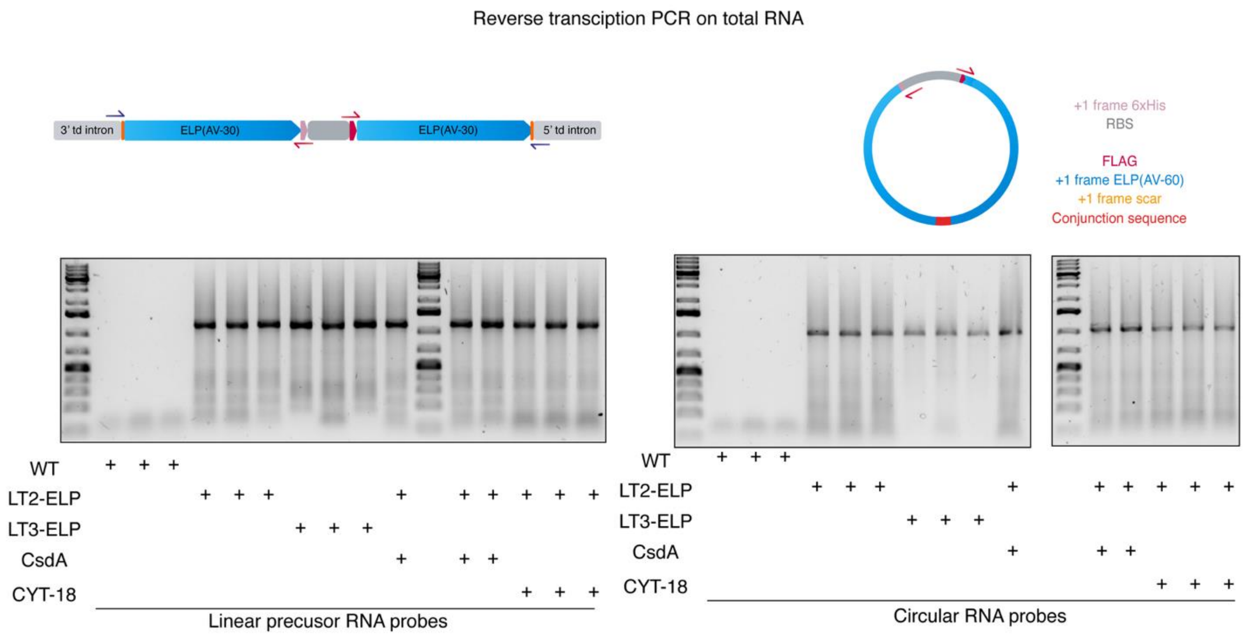

**Supplementary Figure 9**. Reverse transcription to confirm the presence of circular RNA *in vivo*. *E. coli* cells harboring the indicated plasmids were induced to express RNA circularization constructs together with the protein factors CsdA and CYT-18. Total RNA was extracted using TRIzol reagent and reverse-transcribed into cDNA. PCR was performed using primer sets specific for linear precursor RNA or circular RNA to distinguish and validate RNA circularization.

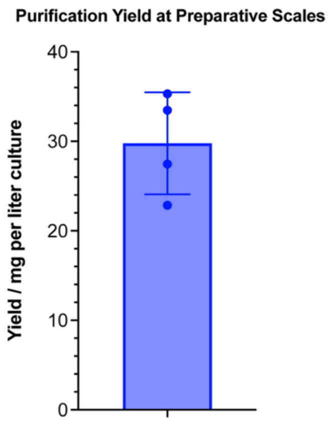

**Supplementary Figure 10**. Purification yield of ELPs synthesized through looped translation at preparative scale. Each data point represents the protein expression ($n$ = 4) with at least a 2-liter shaker-flask culture followed by purification using Ni-NTA agarose resin, as described in the methods.

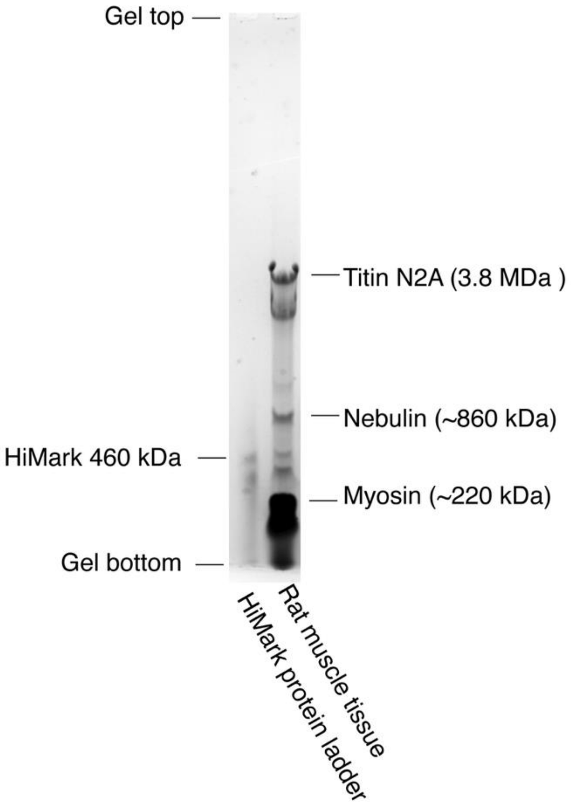

**Supplementary Figure 11**. Protein agarose gel for the analysis of titin molecules extracted from rat muscle tissue using the protocol by Warren and Greaser[7].

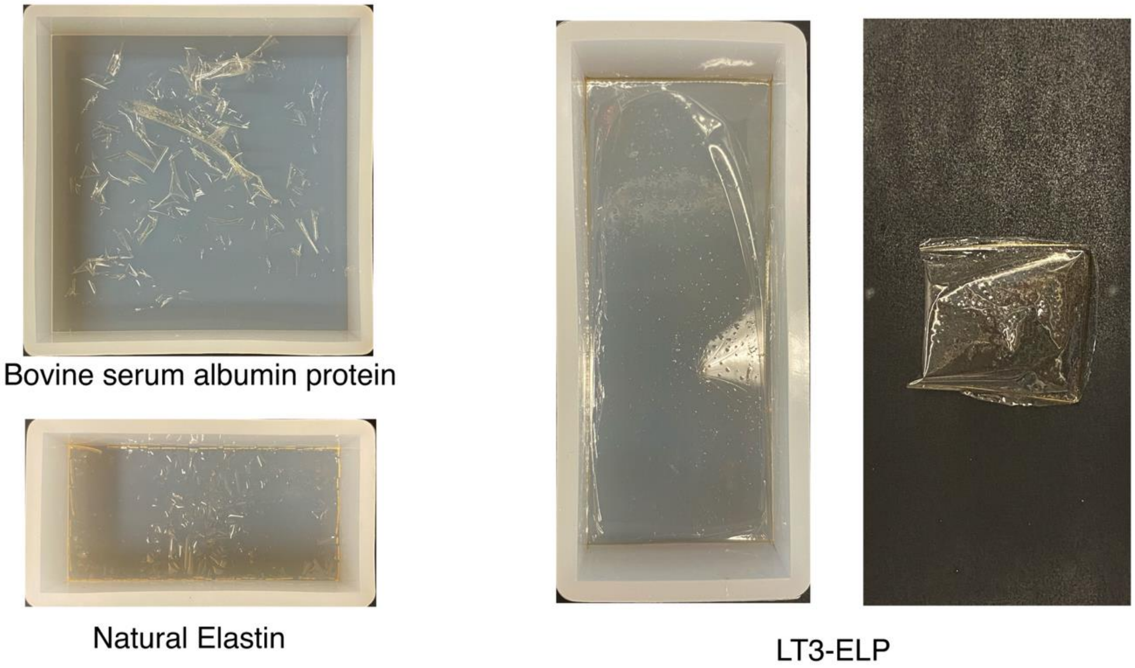

**Supplementary Figure 12**. Fabrication of films composed of different proteins. Briefly: the proteins were dissolved in HFIP solvent at 10% w/v concentration; the solutions were then spread onto a silicone mold, and the solvent was air-dried. Bovine serum albumin (BSA) and natural elastin proteins formed cracked particles, while LT3-ELP formed flexible films without further processing.

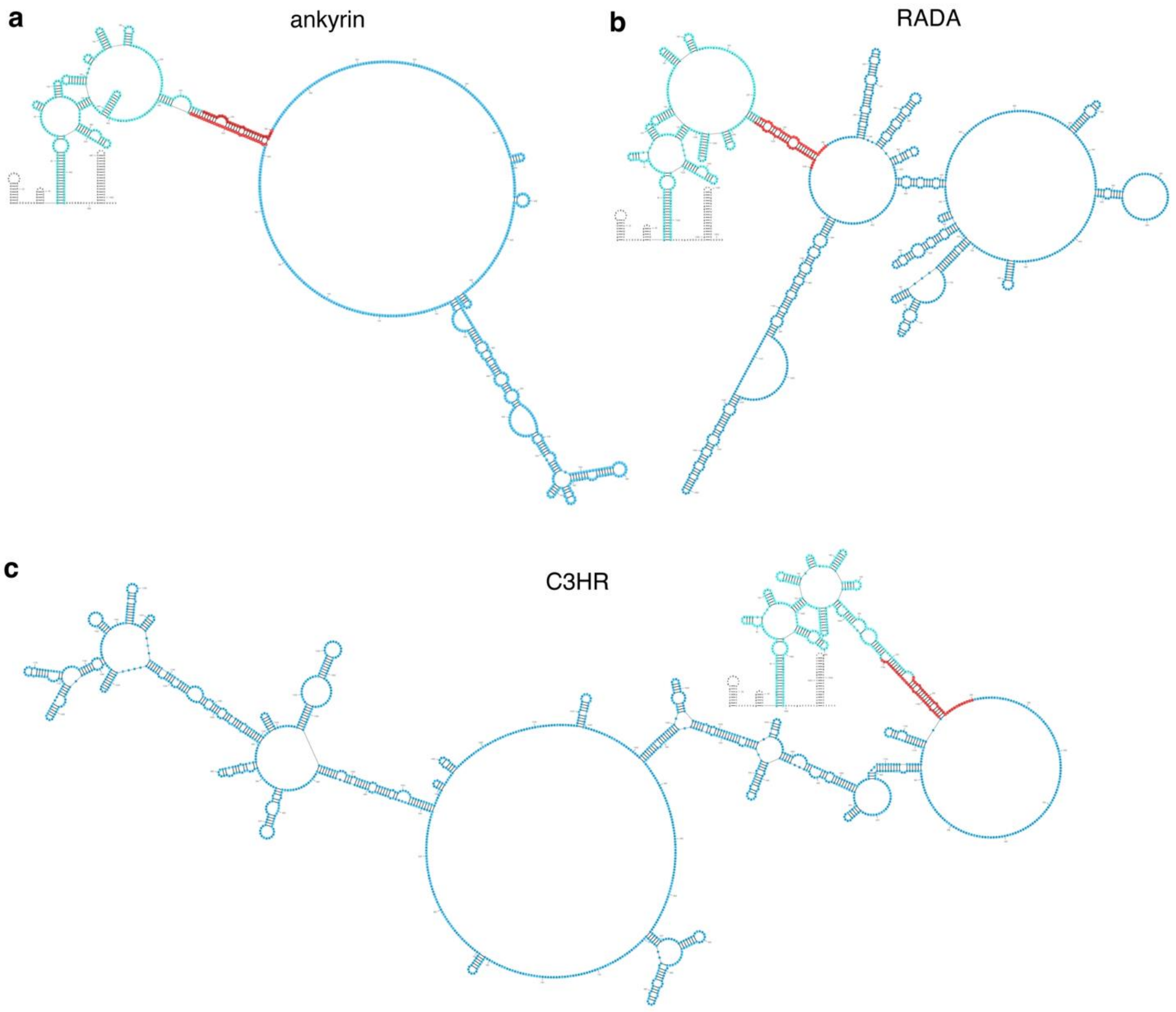
a
ankyrin
b
RADA
c
C3HR

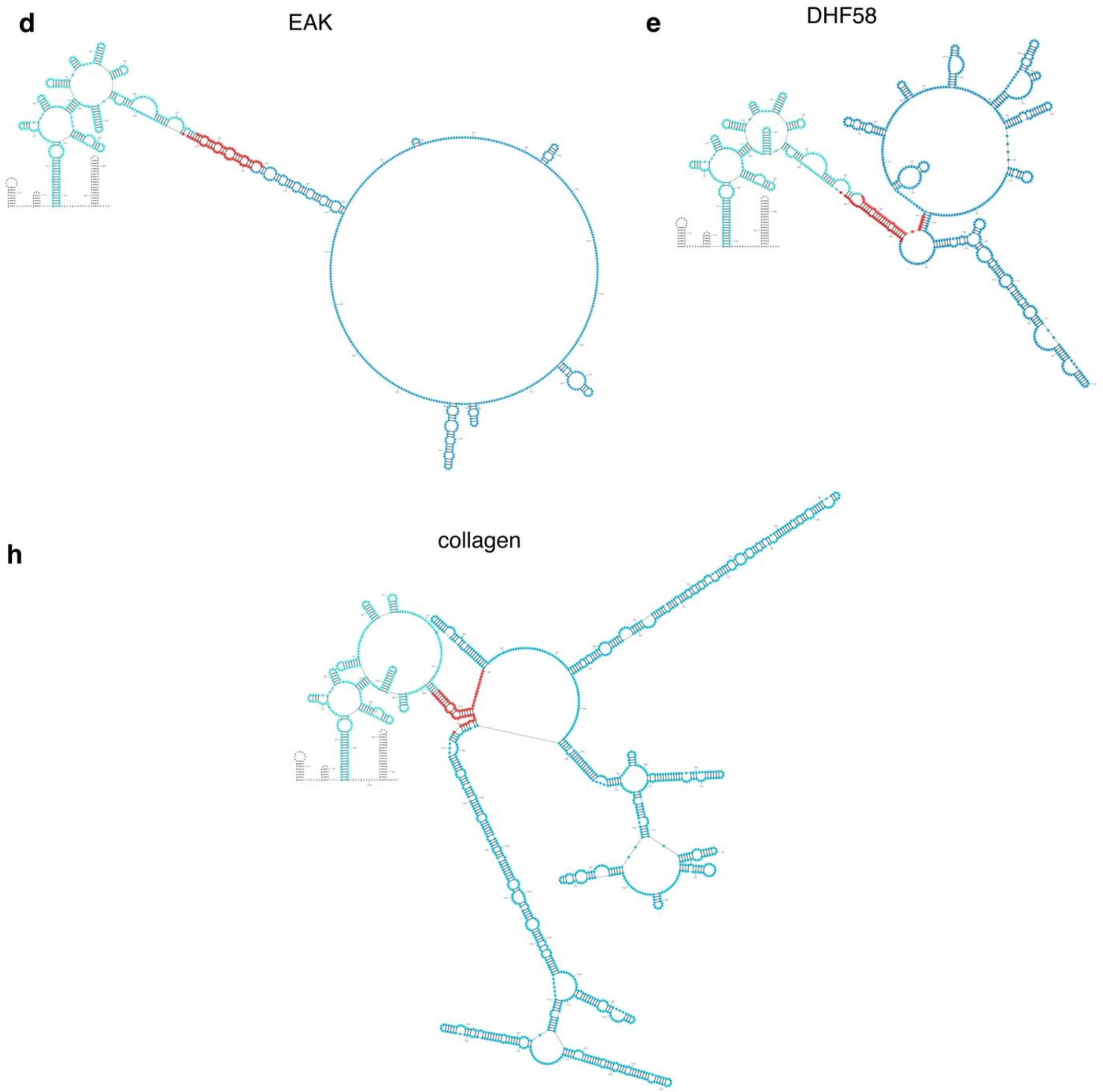

**Supplementary Figure 13.** RNAfold predictions of repetitive proteins: ankyrin (**a**), RADA (**b**), C3HR (**c**), EAK (**d**), DHF58(**e**), and collagen-like protein (**f**) with synonymous locker sequence design. Fibrous protein sequences are colored in blue. Synonymous locker sequences were colored in red. Group **I** intron sequences were colored in cyan. In all cases, the intron folds properly and is insulated from the coding sequences.

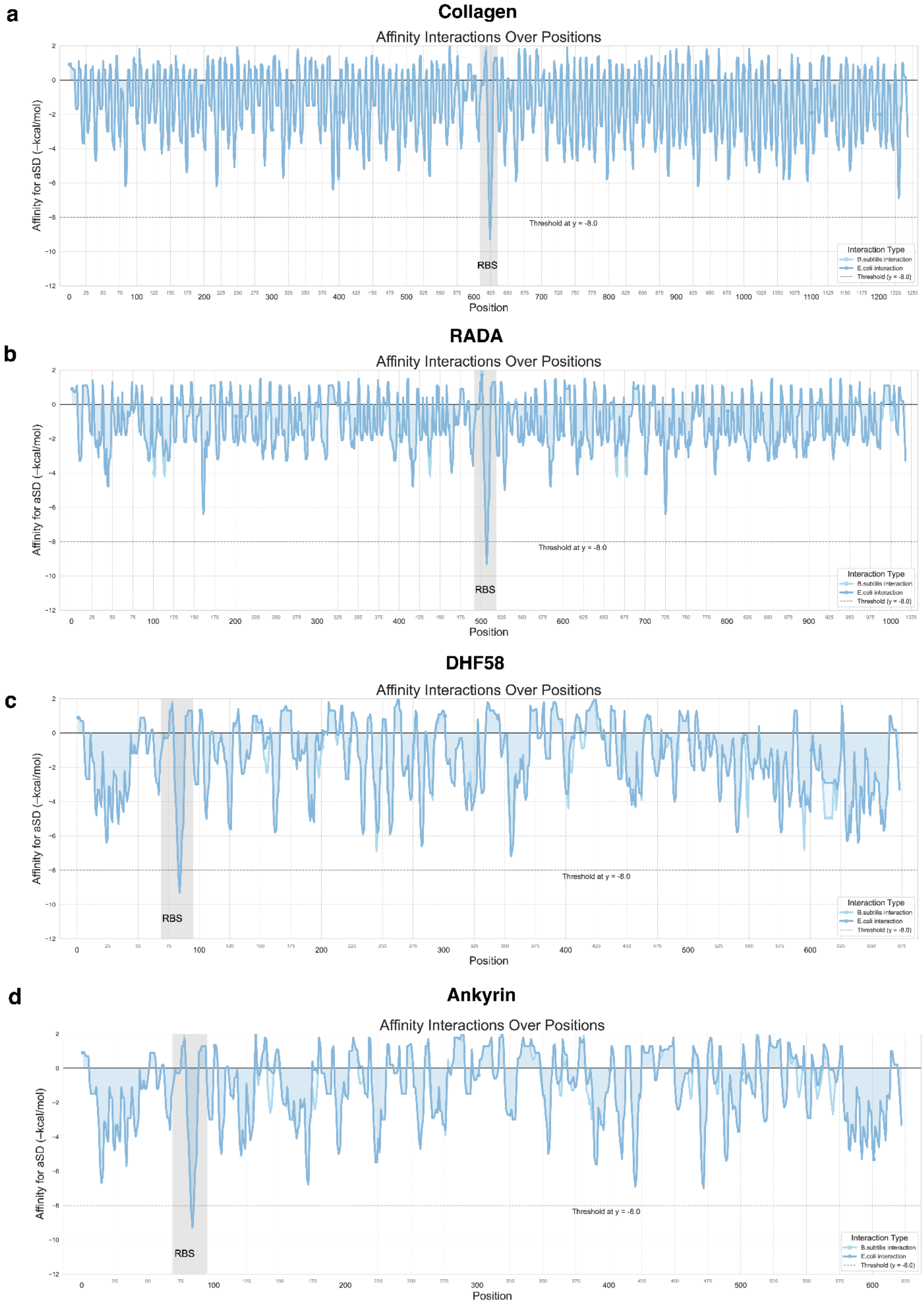
a
Collagen
Affinity Interactions Over Positions
Affinity for aSD (−kcal/mol)
Threshold at y = -8.0
RBS
Interaction Type
B.subtilis interaction
E.coli interaction
Threshold (y = -8.0)
Position
b
RADA
Affinity Interactions Over Positions
Affinity for aSD (−kcal/mol)
Threshold at y = -8.0
RBS
Interaction Type
B.subtilis interaction
E.coli interaction
Threshold (y = -8.0)
Position
c
DHF58
Affinity Interactions Over Positions
Affinity for aSD (−kcal/mol)
Threshold at y = -8.0
RBS
Interaction Type
B.subtilis interaction
E.coli interaction
Threshold (y = -8.0)
Position
d
Ankyrin
Affinity Interactions Over Positions
Affinity for aSD (−kcal/mol)
Threshold at y = -8.0
RBS
Interaction Type
B.subtilis interaction
E.coli interaction
Threshold (y = -8.0)
Position

**Supplementary Figure 14.**

Coding sequence interaction energy calculation probing for collagen (**a**), RADA (**b**), DHF58 (**c**), and ankyrin (**d**). In all cases, the design algorithm described in Fig. S4 and incorporated into LT3-ELP results in the elimination of Shine-Dalgarno-like sequences throughout the coding sequence except for the intended ribosome-binding site.

**a**

tasA$^{SS}$ signal peptide: MGIKKKLSLGVASAALGLALVGGGTWAAFNDIKSKDATFA

tasA$^{SS}$ +1 frame peptide: WVLKRNSVLELPLQHLDWLSLEEEHGQHLTTLNQRMLPLQ

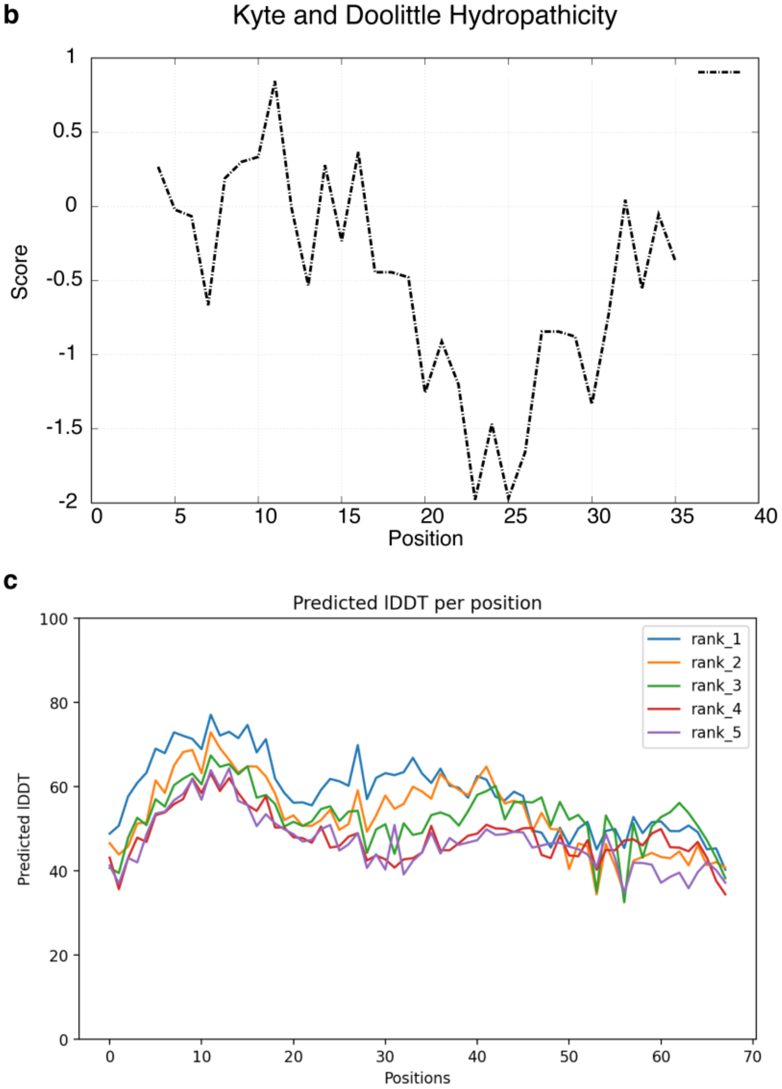

**Supplementary Figure 15**. **a**, Peptide sequence of tasA$^{SS}$ on the 0 frame and +1 frame. **b**, The Kyte-Doolittle hydropathicity[8] calculation of tasA$^{SS}$ +1 frame peptide by the Expasy ProtScale web server[9]. **c**, AlphaFold2 prediction of tasA$^{SS}$ +1 frame peptide by ColabFold[10].

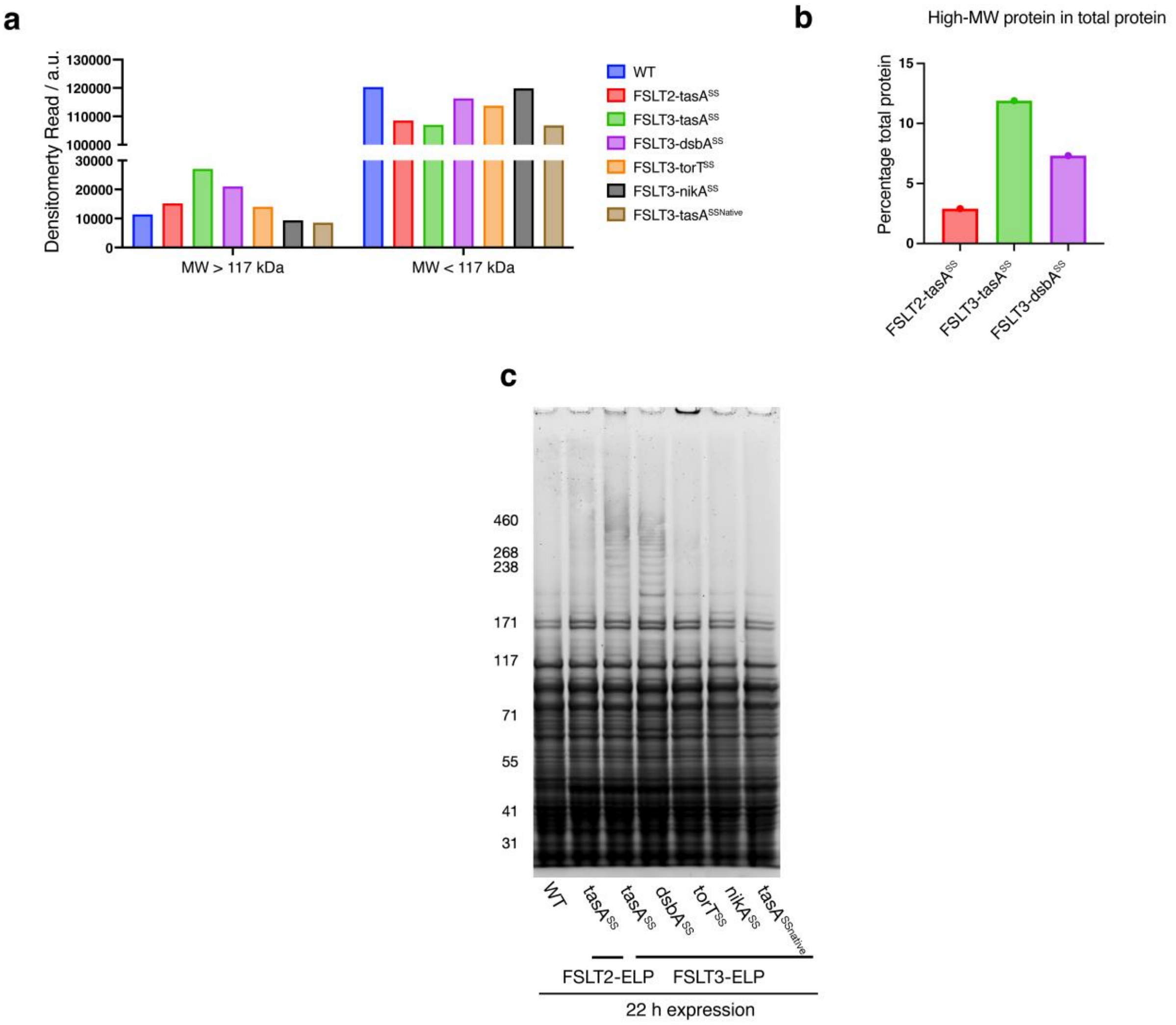

**Supplementary Figure 16**. **a**, Densitometric analysis associated with Figure 5d (in main text) comparing high molecular weight protein to low molecular weight protein. **b**, Fractions of high molecular weight protein relative to total proteins. Calculations are based on the densitometry results in panel **a**. **c**, Total protein gel of expression for 22 hours. Total protein concentrations were quantified using a BCA assay and normalized prior to gel loading.

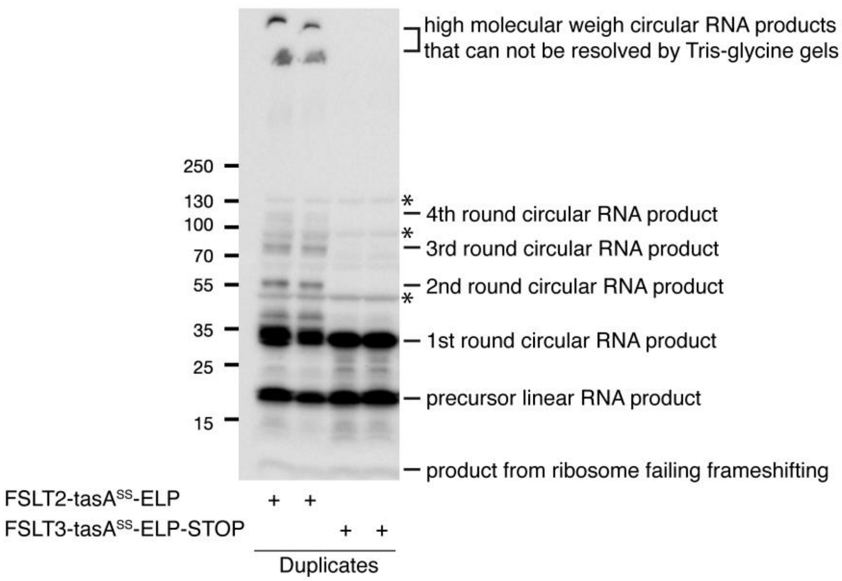

**Supplementary Figure 17**. Evaluation of the RNA circularization efficiency in vivo in the context of a frameshifting looped translator. *E.coli* BL21 cells were used for the expression of FSLT constructs. Cell pellets were collected and lysed using BugBuster Master Mix. Total proteins were separated using a 4-20% Tris-Glycine gel, followed by western blot transfer and detection with anti-FLAG antibodies. Asterisk indicates non-specific detection by anti-FLAG antibodies when using Tris-Glycine gels. These non-specific bands were not detected by anti-FLAG antibodies when using Tris-Acetate gels. Bands corresponding to the n-round of circular RNA translation product were indicated in the figure.

### Supplemental Tables

**Supplemental Table 1. Strains**

| Strain | Description | Sources |
|---|---|---|
| 10-beta | General E. coli cloning strain | New England Biolabs |
| NiCo21(DE3) | General E. coli expression strain | New England Biolabs |
| K12 | E. coli expression strain | Ref: [11] |
| K12ΔsmrB | E. coli expression strain with smrB gene depleted | Ref:[11] |
| K12ΔhrpA | E. coli expression strain with hrpA gene depleted | Ref:[11] |
| K12ΔsmrB ΔhrpA | E. coli expression strain with smrB and hrpA gene depleted | Ref:[11] |
| K07 | Protease deficient strain of B. subtilis WT PY79 (ΔnprE, ΔaprE, Δepr, Δmpr, ΔnprB, Δvpr, Δbpr) | Ref:[12] |
| PG10 | MiniBacillus PG10 | Ref:[1] |

**Supplementary Table 2. Primers used in this study.**

| Primer name | Sequence |
|---|---|
| Pr1 | CCAATTGAATTGGTTCTACATAAATGCCTAACGACTATCCC |
| Pr2 | AGTCCTACAATTTAGCACGGGATTGTC |
| Pr3 | CCGTGCTAAATTGTAGGACTCTCGAGGACGTCCCCGG |
| Pr4 | CATTTATGTAGAACCAATTCAATTGGTGTTGGTTGTTGTTGTGGAATTG |
| Pr5 | GTGCCGGGTACCACCGGTTAATTGAGGCCTGAGTATAAGGTGAC |
| Pr6 | ACCCGGCACTCCTGCGCCGGG |
| Pr7 | GTGGTCCCGGGAACCCCGGCTCCAG |
| Pr8 | AGATCTCGATCCCGCGAAATTAATAC |
| Pr9 | AAAAAAAAACAGATAGGCCCTCCG |
| Pr10 | GGGCCTATCTGTTTTTTTTTCGTCCCC |
| Pr11 | TTGTTATCCGCTCACAATTCCCCTATAGTGAGTCGTATTAATTTCGCGGGATCGA GATCTACGAAAGGGCCTCGTGATACG |
| Pr12 | GCTTACATAAGGAGGAACTACTAATGGACTACAAGGACGACGATGAC |
| Pr13 | CCTCCTTATGTAAGCTTTTTTTTTCGAACTGCGGGTGGC |
| Pr14 | ACTCAATTAAGTGCGGTCGTAACAATGGACTACAAGGACGACGATGAC |
| Pr15 | CGCACTTAATTGAGTGATACCAGATTTTTCGAACTGCGGG |
| Pr16 | CTCAGGCCTCAATTAACCCAAGAAAACATC |
| Pr17 | AACGACCTTATCTGAACATAATGCTACCG |
| Pr18 | TGTTCAGATAAGGTCGTTAATCTTACCCC |
| Pr19 | TTAATTGAGGCCTGAGTATAAGGTGACTTATACTTG |

**Supplementary Table 3. Gene fragments**

| | |
|---|---|
| *LT1-ELP* | GGAATTGTGAGCGGATAACAATTCCCCTCTAGAGAATTGGTTCTACATAAATGCCTAA<br>CGACTATCCCTTTGGGGAGTAGGGTCAAGTGACTCGAAACGATAGACAACTTGCTT<br>TAACAAGTTGGAGATATAGTCTGCTCTGCATGGTGACATGCAGCTGGATATAATTCCG<br>GGGTAAGATTAACGACCTTATCTGAACATAATGCTAGGGGCCGGAGTTCCAGGGGT<br>GGGGGTACCTGGTGCTGGAGTGCCTGGCGTAGGCGTACCGGGTGCTGGCGTGCC<br>AGGCGTGGGAGTGCCCGGAGCCGGAGTCCCAGGGGTAGGTGTGCCTGGGGCTG<br>GTGTACCAGGGGTCGGTGTCCCCGGGGCTGGAGTACCTGGCGTTGGCGTCCCTG<br>GGGCAGGAGTTCCGGGAGTAGGTGTCCCGGGGGCTGGGGTCCCGGGAGTCGGT<br>GTGCCAGGTGCCGGTGTTCCGGGTGTGGGGGTCCCAGGTGCAGGGGTACCCGGG<br>GTTGGCGTTCCAGGAGCCGGCGTTCCGGGCGTTGGTGTCCCTGGCGCGGGAGTT<br>CCTGGGGTGGGTGTGCCGGGAGCTGGAGTCCCGGGTGTAGGCGTGCCTGGTGCA<br>GGAGTACCAGGCGTTGGGGTGCCCGGTGCTGGTGTTCCTGGCGTCGGGGTGCCG<br>GGCTGGCCGCACCATCATCATCACCATTCTGGTATCATTCAATTAAAGGGGGTGTTA<br>ACATGGGTATTAAAAAGAAACTCAGTCTTGGAGTTGCCTCTGCAGCACTTGGATTGG<br>CTCTCGTTGGAGGAGGAACATGGGCAGCATTTAACGACATTAAATCAAAGGATGCTA<br>CCTTTGCAGACTTTAGGGGGTCTCTTTGACCTCGAAGGAGCTGGTGTGCCCGGGG<br>TAGGGGTCCCTGGAGCCGGGGTCCCCGGTGTGGGTGTCCCAGGAGCGGGTGTTC<br>CCGGTGTCGGTGTACCGGGCGCCGGAGTACCCGGTGTAGGTGTTCCAGGCGCTG<br>GAGTTCCCGGAGTCGGGGTTCCAGGTGCTGGGGTTCCGGGGGTGGGAGTCCCTG<br>GTGCGGGGGTTCCTGGTGTCGGAGTGCCAGGAGCTGGGGTACCGGGGGTAGGCG<br>TTCCTGGAGCGGGAGTACCGGGAGTGGGTGTACCTGGAGCAGGCGTCCCAGGCG<br>TAGGTGTACCCGGAGCAGGGGTTCCCGGGGTGGGCGTTCCCGGCGCAGGAGTGC<br>CGGGTGTCGGGGTACCAGGAGCAGGAGTCCCCGGAGTGGGGGTGCCTGGAGCT<br>GGCGTACCAGGTGTCGGCGTACCTGGGGCGGGCGTACCCGGCGTAGGGGTGCCA<br>ACCACCGGTTAATTGAGGCCTGAGTATAAGGTGACTTATACTTGTAATCTATCTAAAC<br>GGGGAACCTCTCTAGTAGACAATCCCGTGCTAAATTGTAGGACTAAACAGATAGGCC<br>CTCTTCGGAGGGCCTATCTGTTTTTTTTT |
| *LT1-ELP*$_{med}$ | GGAATTGTGAGCGGATAACAATTCCCCTCTAGAGAATTGGTTCTACATAAATGCCTAA<br>CGACTATCCCTTTGGGGAGTAGGGTCAAGTGACTCGAAACGATAGACAACTTGCTT<br>TAACAAGTTGGAGATATAGTCTGCTCTGCATGGTGACATGCAGCTGGATATAATTCCG<br>GGGTAAGATTAACGACCTTATCTGAACATAATGCTAGGGGCCGGAGTTCCAGGGGT<br>GGGGGTACCTGGTGCTGGAGTGCCTGGCGTAGGCGTACCGGGTGCTGGCGTGCC<br>AGGCGTGGGAGTGCCCGGAGCCGGAGTCCCAGGGGTAGGTGTGCCTGGGGCTG<br>GTGTACCAGGGGTCGGTGTCCCCGGGGCTGGAGTACCTGGCGTTGGCGTCCCTG<br>GGGCAGGAGTTCCGGGAGTAGGTGTCCCGGGGGCTGGGGTCCCGGGAGTCGGT<br>GTGCCAGGTGCCGGTGTTCCGGGTGTGGGGGTCCCAGGTGCAGGGGTACCCGGG<br>GTTGGCGTTCCAGGAGCCGGCGTTCCGGGCGTTGGTGTCCCTGGCGCGGGAGTT<br>CCTGGGGTGGGTGTGCCGGGAGCTGGAGTCCCGGGTGTAGGCGTGCCTGGTGCA<br>GGAGTACCAGGCGTTGGGGTGCCCGGTGCTGGTGTTCCTGGCGTCGGGGTGCCG<br>GGCTGGCCGCACCATCATCATCACCATTCTGGTATCATTCAATTAAAGGGGGTGTTA<br>ACATGGGTATTAAAAAGAAACTCAGTCTTGGAGTTGCCTCTGCAGCACTTGGATTGG<br>CTCTCGTTGGAGGAGGAACATGGGCAGCATTTAACGACATTAAATCAAAGGATGCTA<br>CCTTTGCAGACTTTAGGGGGTCTCTTTGACCTCGAAGGAGCTGGTGTGCCCGGGG<br>TAGGGGTCCCTGGAGCCGGGGTCCCCGGTGTGGGTGTCCCAGGAGCGGGTGTTC<br>CCGGTGTCGGTGTACCGGGCGCCGGAGTACCCGGTGTAGGTGTTCCAGGCGCTG<br>GAGTTCCCGGAGTCGGGGTTCCAGGTGCTGGGGTTCCGGGGGTGGGAGTCCCTG<br>GTGCGGGGGTTCCTGGTGTCGGAGTGCCAGGAGCTGGGGTACCGGGGGTAGGCG<br>TTCCTGGAGCGGGAGTACCGGGAGTGGGTGTACCTGGAGCAGGCGTCCCAGGCG<br>TAGGTGTACCCGGAGCAGGGGTTCCCGGGGTGGGCGTTCCCGGCGCAGGAGTGC<br>CGGGTACCACCGGTTAATTGAGGCCTGAGTATAAGGTGACTTATACTTGTAATCTATC<br>TAAACGGGGAACCTCTCTAGTAGACAATCCCGTGCTAAATTGTAGGACTAAACAGAT<br>AGGCCCTCTTCGGAGGGCCTATCTGTTTTTTTTT |
| *LT1-ELP*$_{short}$ | GGAATTGTGAGCGGATAACAATTCCCCTCTAGAGAATTGGTTCTACATAAATGCCTAA<br>CGACTATCCCTTTGGGGAGTAGGGTCAAGTGACTCGAAACGATAGACAACTTGCTT |

| | |
|---|---|
| | TAACAAGTTGGAGATATAGTCTGCTCTGCATGGTGACATGCAGCTGGATATAATTCCG<br>GGGTAAGATTAACGACCTTATCTGAACATAATGCTAGGGGCCGGAGTTCCAGGGGT<br>GGGGGTACCTGGTGCTGGAGTGCCTGGCGTAGGCGTACCGGGTGCTGGCGTGCC<br>AGGCGTGGGAGTGCCCGGAGCCGGAGTCCCAGGGGTAGGTGTGCCTGGGGCTG<br>GTGTACCAGGGGTCGGTGTCCCCGGGGCTGGAGTACCTGGCGTTGGCGTCCCTG<br>GGGCAGGAGTTCCGGGAGTAGGTGTCCCGGGGGCTGGGGTCCCGGGAGTCGGT<br>GTGCCAGGTGCCGGTGTTCCGGGTGTGGGGGTCCCAGGTGCAGGGGTACCCGGG<br>GTTGGCGTTCCAGGAGCCGGCGTTCCGGGCGTTGGTGTCCCTGGCGCGGGAGTT<br>CCTGGGGTGGGTGTGCCGGGAGCTGGAGTCCCGGGTGTAGGCGTGCCTGGTGCA<br>GGAGTACCAGGCGTTGGGGTGCCCGGTGCTGGTGTTCCTGGCGTCGGGGTGCCG<br>GGCTGGCCGCACCATCATCATCACCATTCTGGTATCATTCAATTAAAGGGGGTGTTA<br>ACATGGGTATTAAAAAGAAACTCAGTCTTGGAGTTGCCTCTGCAGCACTTGGATTGG<br>CTCTCGTTGGAGGAGGAACATGGGCAGCATTTAACGACATTAAATCAAAGGATGCTA<br>CCTTTGCAGACTTTAGGGGGTCTCTTTGACCTCGAAGGAGCTGGTGTGCCCGGGG<br>TAGGGGTCCCTGGAGCCGGGGTTCCCGGGACCACCGGTTAATTGAGGCCTGAGTA<br>TAAGGTGACTTATACTTGTAATCTATCTAAACGGGGAACCTCTCTAGTAGACAATCCC<br>GTGCTAAATTGTAGGACTAAACAGATAGGCCCTCTTCGGAGGGCCTATCTGTTTTTT<br>TTT |
| *LT2-ELP* | GGAATTGTGAGCGGATAACAATTCCCCTCTAGAGGGAGACCCTCGAATGGAATTGG<br>TTCTACATAAATGCCTAACGACTATCCCTTTGGGGAGTAGGGTCAAGTGACTCGAAA<br>CGATAGACAACTTGCTTTAACAAGTTGGAGATATAGTCTGCTCTGCATGGTGACATG<br>CAGCTGGATATAATTCCGGGGTAAGATTAACGACCTTATCTGAACATAATGCTACCGT<br>TTGGTGCTGGTGTTCCTGGTGCTGGTGTTCCTGGGGCCGGAGTTCCAGGGGTGG<br>GGGTACCTGGTGCTGGAGTGCCTGGCGTAGGCGTACCGGGTGCTGGCGTGCCAG<br>GCGTGGGAGTGCCCGGAGCCGGAGTCCCAGGGGTAGGTGTGCCTGGGGCTGGT<br>GTACCAGGGGTCGGTGTCCCCGGGGCTGGAGTACCTGGCGTTGGCGTCCCTGGG<br>GCAGGAGTTCCGGGAGTAGGTGTCCCGGGGGCTGGGGTCCCGGGAGTCGGTGTG<br>CCAGGTGCCGGTGTTCCGGGTGTGGGGGTCCCAGGTGCAGGGGTACCCGGGGTT<br>GGCGTTCCAGGAGCCGGCGTTCCGGGCGTTGGTGTCCCTGGCGCGGGAGTTCCT<br>GGGGTGGGTGTGCCGGGAGCTGGAGTCCCGGGTGTAGGCGTGCCTGGTGCAGG<br>AGTACCAGGCGTTGGGGTGCCCGGTGCTGGTGTTCCTGGCGTCGGGGTGCCGGG<br>CAGCGCACATCATCACCACCATCATTCTGGTATCATTCAATTAAAGGGGGTGTTAACA<br>ATGGACTACAAGGACGACGATGACAAGGGAGCTGGTGTGCCCGGGGTAGGGGTCC<br>CTGGAGCCGGGGTCCCCGGTGTGGGTGTCCCAGGAGCGGGTGTTCCCGGTGTCG<br>GTGTACCGGGCGCCGGAGTACCCGGTGTAGGTGTTCCAGGCGCTGGAGTTCCCG<br>GAGTCGGGGTTCCAGGTGCTGGGGTTCCGGGGGTGGGAGTCCCTGGTGCGGGG<br>GTTCCTGGTGTCGGAGTGCCAGGAGCTGGGGTACCGGGGGTAGGCGTTCCTGGA<br>GCGGGAGTACCGGGAGTGGGTGTACCTGGAGCAGGCGTCCCAGGCGTAGGTGTA<br>CCCGGAGCAGGGGTTCCCGGGGTGGGCGTTCCCGGCGCAGGAGTGCCGGGTGT<br>CGGGGTACCAGGAGCAGGAGTCCCCGGAGTGGGGGTGCCTGGAGCTGGCGTACC<br>AGGTGTCGGCGTACCTGGGGCGGGCGTACCCGGCGTAGGGGTGCCAGGTGCTGG<br>AGTACCAGGAGCAGGTGTACCAGATGTTTTCTTGGGTTAATTGAGGCCTGAGTATAA<br>GGTGACTTATACTTGTAATCTATCTAAACGGGGAACCTCTCTAGTAGACAATCCCGTG<br>CTAAATTGTAGGACTAATTCCATTTATCAGATTTCTAGAAACAGATAGGCCCTCTTCG<br>GAGGGCCTATCTGTTTTTTTTT |
| *RBSstr* | GGAATTGTGAGCGGATAACAATTCCCCTCTAGAGGGAGACCCTCGAATGGAATTGG<br>TTCTACATAAATGCCTAACGACTATCCCTTTGGGGAGTAGGGTCAAGTGACTCGAAA<br>CGATAGACAACTTGCTTTAACAAGTTGGAGATATAGTCTGCTCTGCATGGTGACATG<br>CAGCTGGATATAATTCCGGGGTAAGATTAACGACCTTATCTGAACATAATGCTACCGT<br>TTGGTGCTGGTGTTCCTGGTGCTGGTGTTCCTGGGGCCGGAGTTCCAGGGGTGG<br>GGGTACCTGGTGCTGGAGTGCCTGGCGTAGGCGTACCGGGTGCTGGCGTGCCAG<br>GCGTGGGAGTGCCCGGAGCCGGAGTCCCAGGGGTAGGTGTGCCTGGGGCTGGT<br>GTACCAGGGGTCGGTGTCCCCGGGGCTGGAGTACCTGGCGTTGGCGTCCCTGGG<br>GCAGGAGTTCCGGGAGTAGGTGTCCCGGGGGCTGGGGTCCCGGGAGTCGGTGTG<br>CCAGGTGCCGGTGTTCCGGGTGTGGGGGTCCCAGGTGCAGGGGTACCCGGGGTT<br>GGCGTTCCAGGAGCCGGCGTTCCGGGCGTTGGTGTCCCTGGCGCGGGAGTTCCT |

| | |
|---|---|
| | GGGGTGGGTGTGCCGGGAGCTGGAGTCCCGGGTGTAGGCGTGCCTGGTGCAGG<br>AGTACCAGGCGTTGGGGTGCCCGGTGCTGGTGTTCCTGGCGTCGGGGTGCCGGG<br>CAGCGCATGGAGCCACCCGCAGTTCGAAAAAAAAAGCTTACATAAGGAGGAACTAC<br>TAATGGACTACAAGGACGACGATGACAAGGGAGCTGGTGTGCCCGGGGTAGGGGT<br>CCCTGGAGCCGGGGTCCCCGGTGTGGGTGTCCCAGGAGCGGGTGTTCCCGGTGT<br>CGGTGTACCGGGCGCCGGAGTACCCGGTGTAGGTGTTCCAGGCGCTGGAGTTCC<br>CGGAGTCGGGGTTCCAGGTGCTGGGGTTCCGGGGGTGGGAGTCCCTGGTGCGG<br>GGGTTCCTGGTGTCGGAGTGCCAGGAGCTGGGGTACCGGGGGTAGGCGTTCCTG<br>GAGCGGGAGTACCGGGAGTGGGTGTACCTGGAGCAGGCGTCCCAGGCGTAGGTG<br>TACCCGGAGCAGGGGTTCCCGGGGTGGGCGTTCCCGGCGCAGGAGTGCCGGGT<br>GTCGGGGTACCAGGAGCAGGAGTCCCCGGAGTGGGGGTGCCTGGAGCTGGCGTA<br>CCAGGTGTCGGCGTACCTGGGGCGGGCGTACCCGGCGTAGGGGTGCCAGGTGCT<br>GGAGTACCAGGAGCAGGTGTACCAGATGTTTTCTTGGGTTAATTGAGGCCTGAGTAT<br>AAGGTGACTTATACTTGTAATCTATCTAAACGGGGAACCTCTCTAGTAGACAATCCCG<br>TGCTAAATTGTAGGACTAATTCCATTTATCAGATTTCTAGAAACAGATAGGCCCTCTTC<br>GGAGGGCCTATCTGTTTTTTTTT |
| *RBSweak* | GGAATTGTGAGCGGATAACAATTCCCCTCTAGAGGGAGACCCTCGAATGGAATTGG<br>TTCTACATAAATGCCTAACGACTATCCCTTTGGGGAGTAGGGTCAAGTGACTCGAAA<br>CGATAGACAACTTGCTTTAACAAGTTGGAGATATAGTCTGCTCTGCATGGTGACATG<br>CAGCTGGATATAATTCCGGGGTAAGATTAACGACCTTATCTGAACATAATGCTACCGT<br>TTGGTGCTGGTGTTCCTGGTGCTGGTGTTCCTGGGGCCGGAGTTCCAGGGGTGG<br>GGGTACCTGGTGCTGGAGTGCCTGGCGTAGGCGTACCGGGTGCTGGCGTGCCAG<br>GCGTGGGAGTGCCCGGAGCCGGAGTCCCAGGGGTAGGTGTGCCTGGGGCTGGT<br>GTACCAGGGGTCGGTGTCCCCGGGGCTGGAGTACCTGGCGTTGGCGTCCCTGGG<br>GCAGGAGTTCCGGGAGTAGGTGTCCCGGGGGCTGGGGTCCCGGGAGTCGGTGTG<br>CCAGGTGCCGGTGTTCCGGGTGTGGGGGTCCCAGGTGCAGGGGTACCCGGGGTT<br>GGCGTTCCAGGAGCCGGCGTTCCGGGCGTTGGTGTCCCTGGCGCGGGAGTTCCT<br>GGGGTGGGTGTGCCGGGAGCTGGAGTCCCGGGTGTAGGCGTGCCTGGTGCAGG<br>AGTACCAGGCGTTGGGGTGCCCGGTGCTGGTGTTCCTGGCGTCGGGGTGCCGGG<br>CAGCGCATGGAGCCACCCGCAGTTCGAAAAATCTGGTATCACTCAATTAAGTGCGG<br>TCGTAACAATGGACTACAAGGACGACGATGACAAGGGAGCTGGTGTGCCCGGGGT<br>AGGGGTCCCTGGAGCCGGGGTCCCCGGTGTGGGTGTCCCAGGAGCGGGTGTTCC<br>CGGTGTCGGTGTACCGGGCGCCGGAGTACCCGGTGTAGGTGTTCCAGGCGCTGG<br>AGTTCCCGGAGTCGGGGTTCCAGGTGCTGGGGTTCCGGGGGTGGGAGTCCCTGG<br>TGCGGGGGTTCCTGGTGTCGGAGTGCCAGGAGCTGGGGTACCGGGGGTAGGCGT<br>TCCTGGAGCGGGAGTACCGGGAGTGGGTGTACCTGGAGCAGGCGTCCCAGGCGT<br>AGGTGTACCCGGAGCAGGGGTTCCCGGGGTGGGCGTTCCCGGCGCAGGAGTGCC<br>GGGTGTCGGGGTACCAGGAGCAGGAGTCCCCGGAGTGGGGGTGCCTGGAGCTG<br>GCGTACCAGGTGTCGGCGTACCTGGGGCGGGCGTACCCGGCGTAGGGGTGCCAG<br>GTGCTGGAGTACCAGGAGCAGGTGTACCAGATGTTTTCTTGGGTTAATTGAGGCCT<br>GAGTATAAGGTGACTTATACTTGTAATCTATCTAAACGGGGAACCTCTCTAGTAGACA<br>ATCCCGTGCTAAATTGTAGGACTAATTCCATTTATCAGATTTCTAGAAACAGATAGGC<br>CCTCTTCGGAGGGCCTATCTGTTTTTTTTT |
| *ΔSD (LT3-ELP)* | GGAATTGTGAGCGGATAACAATTCCCCTCTAGAGGGAGACCCTCGAATGGAATTGG<br>TTCTACATAAATGCCTAACGACTATCCCTTTGGGGAGTAGGGTCAAGTGACTCGAAA<br>CGATAGACAACTTGCTTTAACAAGTTGGAGATATAGTCTGCTCTGCATGGTGACATG<br>CAGCTGGATATAATTCCGGGGTAAGATTAACGACCTTATCTGAACATAATGCTACCGT<br>TTGTAGGCGTTCCGGGTGCAGGGGTGCCGGGTGGAGCCGGCGTCCCGGGCGTTG<br>GGGTCCCTGGGGCGGGAGTTCCCGGTGTCGGCGTTCCAGGAGCCGGAGTTCCTG<br>GCGTAGGTGTCCCAGGAGCTGGGGTGCCCGGGGTAGGCGTACCCGGGGCCGGA<br>GTCCCGGGTGTTGGCGTGCCCGGTGCTGGGGTTCCCGGCGTAGGCGTGCCGGGT<br>GCAGGAGTCCCTGGCGTTGGCGTACCAGGAGCGGGAGTCCCAGGGGTGGGCGTT<br>CCCGGAGCAGGAGTACCCGGAGTGGGTGTGCCTGGAGCCGGGGTGCCTGGTGTC<br>GGTGTCCCTGGAGCAGGCGTCCCAGGCGTGGGTGTCCCGGGAGCAGGGGTACCC<br>GGCGTTGGTGTCCCCGGAGCTGGCGTCCCCGGTGTGGGTGTTCCTGGAGCGGGT<br>GTACCGGGAGTCGGTGTACCTGGTGCAGGTGTGCCGGGAGTGGGCGTCCCTTCG |

| | |
|---|---|
| | GGACATCATCACCACCATCATTCTGGTATCATTCAATTAAAGGGGGTGTTAACAATGG<br>ACTACAAGGACGACGATGACAAGGGTGCGGGAGTACCGGGTGTCGGGGTTCCTGG<br>GGCAGGTGTACCCGGTGTAGGCGTTCCTGGTGCTGGAGTCCCCGGGGTCGGTGT<br>GCCAGGTGCTGGTGTGCCCGGAGTCGGCGTACCTGGAGCTGGAGTTCCGGGTGT<br>GGGCGTACCGGGCGCTGGGGTACCTGGCGTGGGAGTGCCAGGCGCAGGAGTGC<br>CTGGGGTCGGGGTACCAGGTGCCGGAGTACCAGGCGTAGGAGTTCCAGGGGCTG<br>GAGTGCCGGGCGTCGGGGTCCCAGGTGCAGGGGTTCCAGGCGTTGGAGTACCTG<br>GGGCTGGTGTTCCCGGGGTTGGCGTTCCGGGAGCTGGTGTACCAGGGGTAGGGG<br>TCCCCGGCGCGGGCGTGCCAGGGGTCGGCGTGCCTGGCGCCGGTGTTCCGGGC<br>GTAGGGGTGCCAGGAGCAGGTGTTCCAGGTGTCGGAGTGCCCGTCGGCGTACCC<br>GGTGCCGGCGTACCTGGCGATGTTTTCTTGGGTTAATTGAGGCCTGAGTATAAGGT<br>GACTTATACTTGTAATCTATCTAAACGGGGAACCTCTCTAGTAGACAATCCCGTGCTA<br>AATTGTAGGACTAATTCCATTTATCAGATTTCTAGAAACAGATAGGCCCTCTTCGGAG<br>GGCCTATCTGTTTTTTTTT |
| *DHF58* | GGAATTGTGAGCGGATAACAATTCCCCTCTAGAGGGAGACCCTCGAATGGAATTGG<br>TTCTACATAAATGCCTAACGACTATCCCTTTGGGGAGTAGGGTCAAGTGACTCGAAA<br>CGATAGACAACTTGCTTTAACAAGTTGGAGATATAGTCTGCTCTGCATGGTGACATG<br>CAGCTGGATATAATTCCGGGGTAAGATTAACGACCTTATCTGAACATAATGCTACCGT<br>TTGGTTCCGGCGGATCCGGTGGATCGGGCGGGTCGGGACATCATCACCACCATCA<br>TTCTGGTATCATTCAATTAAAGGGGGTGTTAACAATGGACTACAAGGACGACGATGA<br>CAAGATGGAGCTCCTTAGAGTCGTCATGTTAGTAAAAGAAGCAGAGGAATTGCTTAC<br>GCTGGCAGTCATCAAAGGATCAGAAGACGACTTACAAAAAGCTTTGAGAACAGCTG<br>TCGAGGCCGCACGAGAGGCAGTTAAGGTGCTTTTACAGGCAGTGAAACGCGGAGA<br>CCCGGAAGTAGCATTACGTGCGGTTGAACTCGTGGTAAGGGTAGCGGAATTACTGC<br>TGCGCATCGCGAAGGAGAGTGGCAGTGAGCTTGCCCTCAAAATGGCCTTACTTGTA<br>GCGGAAGAAGCAGCCAGACTTGCTAAAATAGTTCTTGAGCTTGCTGAGAAACAGGG<br>CGATCCGGAAGTTGCGCGTCGGGCCGTCGAACTCGTGAAACGCGTTGCTGAGCTG<br>CTCGAGCGAATCGCTCGTGAATCAGGCTCAGAGGAAGCGAAAGAGCGTGCAGAGC<br>GTGTTCGTGAAGAGGCGCGGGAATTACAGGAAAGGGTCAAAGAGCTGAGAGAGAG<br>AGAAGGCTTAGAGGGATCCGGTGGATCTGGCGGAAGTGGCGGGGATGTTTTCTTG<br>GGTTAATTGAGGCCTGAGTATAAGGTGACTTATACTTGTAATCTATCTAAACGGGGAA<br>CCTCTCTAGTAGACAATCCCGTGCTAAATTGTAGGACTAATTCCATTTATCAGATTTCT<br>AGAAACAGATAGGCCCTCTTCGGAGGGCCTATCTGTTTTTTTTT |
| *collagen-like protein* | GGAATTGTGAGCGGATAACAATTCCCCTCTAGAGGGAGACCCTCGAATGGAATTGG<br>TTCTACATAAATGCCTAACGACTATCCCTTTGGGGAGTAGGGTCAAGTGACTCGAAA<br>CGATAGACAACTTGCTTTAACAAGTTGGAGATATAGTCTGCTCTGCATGGTGACATG<br>CAGCTGGATATAATTCCGGGGTAAGATTAACGACCTTATCTGAACATAATGCTACCGT<br>TTCCCGGCACTCCAGGTCCACAGGGCCTCCCTGGGGCTCCCGGCACTCCTGGAC<br>CACAAGGTCTACCTGGAAGTCCGGGAGCTCCTGGCACACCAGGCCCCCAAGGTCT<br>TCCAGGCTCGCCAGGCGCACCAGGAACTCCTGGTCCGCAAGGCCTTCCTGGGTC<br>CCCCGGTGCTCCTGGAACTCCAGGGCCTCAAGGTCTGCCTGGCAGCCCGGGAGC<br>ACCTGGTACACCAGGACCGCAAGGATTACCTGGTTCACCCGGCGCTCCCGGTACT<br>CCTGGCCCACAAGGACTCCCTGGTAGCCCTGGCGCTCCGGGGACTCCAGGACCT<br>CAAGGCTTACCAGGTTCACCTGGCGCACCTGGCACTCCCGGTCCACAAGGCTTGC<br>CTGGGAGTCCCGGAGCTCCAGGCACGCCAGGACCACAGGGTCTTCCTGGATCAC<br>CTGGGGCCCCTGGCACCCCAGGTCCACAGGGACTTCCCGGTTCACCAGGGGCTC<br>CTGGGACTCCTGGGCCGCAAGGGTTACCTGGATCTCCTGGAGCACCAGGCACTCC<br>AGGCCCGCAAGGTTTACCAGGCTCACCTTCGGGACATCATCACCACCATCATTCTG<br>GTATCATTCAATTAAAGGGGGTGTTAACAATGGACTACAAGGACGACGATGACAAGG<br>GTGCCCCCGGAACCCCCGGCCCTCAGGGCTTACCTGGCTCTCCTGGGGCTCCGG<br>GCACTCCGGGTCCGCAGGGATTACCAGGGTCGCCTGGGGCACCTGGGACACCTG<br>GTCCCCAGGGATTGCCTGGTTCTCCAGGTGCACCAGGGACACCAGGGCCACAAG<br>GGCTTCCAGGATCACCAGGCGCTCCAGGTACACCGGGCCCTCAAGGGTTGCCAG<br>GTTCCCCTGGAGCCCCGGGTACCCCAGGACCCCAGGGTTTACCTGGGTCACCAGG<br>AGCGCCCGGTACCCCTGGACCTCAGGGACTACCGGGTTCTCCGGGTGCTCCGGG<br>TACACCTGGACCCCAAGGACTACCAGGATCTCCCGGTGCACCCGGGACCCCTGGT |

| | |
|---|---|
| | CCTCAAGGACTTCCGGGATCACCGGGTGCCCCGGGAACACCAGGTCCTCAGGGG<br>TTACCAGGAAGTCCAGGCGCCCCTGGAACACCTGGGCCTCAGGGTCTACCAGGTA<br>GCCCAGGGGCACCGGGTACTCCAGGTCCCCAAGGCCTCCCTGGTTCACCTGGGA<br>CGCCAGGACCTCAAGGTCTGCCGGGAGATGTTTTCTTGGGTTAATTGAGGCCTGAG<br>TATAAGGTGACTTATACTTGTAATCTATCTAAACGGGGAACCTCTCTAGTAGACAATCC<br>CGTGCTAAATTGTAGGACTAATTCCATTTATCAGATTTCTAGAAACAGATAGGCCCTC<br>TTCGGAGGGCCTATCTGTTTTTTTTT |
| *EAK* | GGAATTGTGAGCGGATAACAATTCCCCTCTAGAGGGAGACCCTCGAATGGAATTGG<br>TTCTACATAAATGCCTAACGACTATCCCTTTGGGGAGTAGGGTCAAGTGACTCGAAA<br>CGATAGACAACTTGCTTTAACAAGTTGGAGATATAGTCTGCTCTGCATGGTGACATG<br>CAGCTGGATATAATTCCGGGGTAAGATTAACGACCTTATCTGAACATAATGCTACCGT<br>TTGCCGAAGCTAAGGCGAAGGCCGAGGCAGAGGCAGAGGCTGAAGCTAAGGCTAA<br>GGCTGAGGCAGAAGCAAAGGCGAAAGCTGAAGCGGAAGCTAAAGCAAAAGCGGA<br>AGCAGAAGCTAAAGCGAAAGCAGAAGCAGAAGCCAAAGCAAAGGCTGAAGCCGAA<br>GCGAAAGCTAAGGCAGAAGCCGAGGCAAAAGCAAAAGCAGAAGCGGAGGCGAAG<br>GCAAAAGCTGAGGCGGAGGCTAAAGCCAAATCGGGACATCATCACCACCATCATTC<br>TGGTATCATTCAATTAAAGGGGGTGTTAACAATGGACTACAAGGACGACGATGACAA<br>GGCTGAAGCAGAGGCTAAAGCTAAAGCAGAGGCAGAAGCTAAGGCCAAGGCAGAG<br>GCCGAGGCTAAGGCAAAGGCCGAAGCAGAAGCAAAAGCTAAAGCTGAAGCTGAAG<br>CCAAGGCTAAAGCTGAGGCTGAAGCGAAGGCTAAAGCAGAAGCTGAAGCTAAAGC<br>TAAGGCGGAAGCTGAGGCCAAAGCTAAAGCCGAAGCTGAAGCAAAAGCCAAGGCC<br>GAGGCCAAAGCCAAAGCTGAGGCGGAGGATGTTTTCTTGGGTTAATTGAGGCCTGA<br>GTATAAGGTGACTTATACTTGTAATCTATCTAAACGGGGAACCTCTCTAGTAGACAAT<br>CCCGTGCTAAATTGTAGGACTAATTCCATTTATCAGATTTCTAGAAACAGATAGGCCC<br>TCTTCGGAGGGCCTATCTGTTTTTTTTT |
| *RADA* | GGAATTGTGAGCGGATAACAATTCCCCTCTAGAGGGAGACCCTCGAATGGAATTGG<br>TTCTACATAAATGCCTAACGACTATCCCTTTGGGGAGTAGGGTCAAGTGACTCGAAA<br>CGATAGACAACTTGCTTTAACAAGTTGGAGATATAGTCTGCTCTGCATGGTGACATG<br>CAGCTGGATATAATTCCGGGGTAAGATTAACGACCTTATCTGAACATAATGCTACCGT<br>TTGCGGATGCACGCGCCGATGCCCGGGCTGATGCAAGGGCGGACGCTCGTGCAG<br>ATGCTCGTGCTGACGCTCGCGCCGATGCAAGAGCGGATGCGAGAGCAGATGCGAG<br>AGCGGATGCACGAGCCGACGCGCGAGCTGACGCTAGAGCTGACGCGAGGGCTGA<br>TGCACGCGCGGACGCTAGAGCCGACGCCCGAGCTGATGCAAGAGCTGACGCAAG<br>AGCTGATGCTAGGGCAGACGCTAGGGCAGATGCACGTGCGGATGCAAGAGCCGAT<br>GCTCGTGCGGACGCAAGAGCAGACGCACGAGCGGACGCACGCGCCGACGCTAGA<br>GCAGACGCTCGGGCAGATGCTAGAGCAGATGCAAGAGCAGATGCCCGTGCAGACG<br>CTAGAGCGGATGCTCGAGCCGATGCTAGGGCGGATGCCAGAGCCGATGCGAGAGC<br>CGATGCACGTGCAGATGCATCGGGACATCATCACCACCATCATTCTGGTATCATTCA<br>ATTAAAGGGGGTGTTAACAATGGACTACAAGGACGACGATGACAAGCGAGCTGATG<br>CTCGTGCCGATGCCAGAGCTGATGCCCGGGCGGATGCTAGAGCCGATGCCCGCG<br>CTGATGCAAGGGCTGATGCTAGAGCGGACGCCCGTGCTGATGCACGAGCAGACGC<br>GAGAGCTGATGCGAGAGCTGACGCCCGCGCAGATGCAAGGGCCGACGCACGTGC<br>TGATGCGAGGGCAGATGCCAGGGCAGATGCTCGGGCTGATGCTCGCGCTGACGCA<br>CGGGCCGATGCAAGGGCAGATGCGCGTGCTGATGCTCGAGCAGATGCCAGAGCG<br>GATGCCCGAGCAGATGCTCGAGCGGATGCTAGGGCTGATGCGCGAGCAGATGCAC<br>GGGCTGACGCCAGGGCTGATGCCAGAGCAGACGCCAGAGCAGATGCTAGGGCCG<br>ATGCTAGAGCTGATGCACGTGCCGACGCTCGAGCTGATGCCGATGCTCGGGCAGA<br>CGCGCGCGCAGACGCAGATGTTTTCTTGGGTTAATTGAGGCCTGAGTATAAGGTGA<br>CTTATACTTGTAATCTATCTAAACGGGGAACCTCTCTAGTAGACAATCCCGTGCTAAA<br>TTGTAGGACTAATTCCATTTATCAGATTTCTAGAAACAGATAGGCCCTCTTCGGAGGG<br>CCTATCTGTTTTTTTTT |
| *C3HR* | GGAATTGTGAGCGGATAACAATTCCCCTCTAGAGGGAGACCCTCGAATGGAATTGG<br>TTCTACATAAATGCCTAACGACTATCCCTTTGGGGAGTAGGGTCAAGTGACTCGAAA<br>CGATAGACAACTTGCTTTAACAAGTTGGAGATATAGTCTGCTCTGCATGGTGACATG<br>CAGCTGGATATAATTCCGGGGTAAGATTAACGACCTTATCTGAACATAATGCTACCGT<br>TTTCCGGCGGATCCGGCGGATCGGGTGGATCATCGGGACATCATCACCACCATCAT |

| | |
|---|---|
| | TCTGGTATCATTCAATTAAAGGGGGTGTTAACAATGGACTACAAGGACGACGATGAC<br>AAGATGGTATATGAAAGACTTAAGGAATTAATTGAAGAAAATCCGGAGATTCAAGAGA<br>TTCTCGAGTTATGGAAATTTATTGGACGCGAAGACGTCGTGGAAAAACTTTTTGAGG<br>TCATAAAATGGGCTGTCGAGGAGGGAGGCAATAATATGGTTTTCATCTCTCTTTTAAA<br>AGAAATTTTATCCAACCCTGAGATTTTTGAAATTTTATTGAAATGGGTCGACATCGGC<br>CGCGAAGACGTAGTGAAAAAACTTTTTTCTGTAATCAAACAGGCAGTAGAGGACGG<br>AGGGAATAACATGGTCTTTCTTTCTCTCTTGAAGGAACTCCTGAGCAATCCGGAAAT<br>CTTTGAAATACTGCTGTTGTGGGTAGAGATTAATCGAGAAGACGTCGTAAAAAAGTT<br>TTTTGATGTAATTAAATGGGCTGTCGAAAGAGGCGGCAACAACCCTAAGTTTCTTTC<br>GTTATTGAAATTGATCCTTTCTAATCCCGAAATATTTAAAATTTTATTACTGTGGGTTTA<br>TGAGAACAAGGAGGAGGTAGTTAAGAAACTCTTTGATGTTTTGCTTTACGCGGTCAA<br>AGAAGGCGGGGACAACGAAAAGTTCTTGGAATTACTGAAGGAAATCCTCAGCAATC<br>CGGAGATTTTTGAGATTCTTTTGGAGTGGGTCGAAGAAAATAAGGAAGATGTAGTGA<br>AAAAATTATTTGATGTGATCAAATATGCAGTAGAGCGGGGCGGTGATAATGAAAAATT<br>TTATGAACTGCTGAAACTTCTCCTGTCGAATCCGGAGATCTTCAAGATTCTGCTCGA<br>GTGGGTTGAAATTAACAAAGAAGACGTGGTTAAAAAATTTTTTGATGTCATTAAATGG<br>GCTGTCGAAGAGGGGGGTGACAACCCCGTATTTTATCGTCTCCTTAAACTTATTCTG<br>TCCAACCCGGAGATTTTTAAAATTCTGCTCGAATGGGTGGATATCAACAATGAAGATG<br>TTGTGAAAAAATTATTTTCCGTAATCAAACAGGCAGTGGAAGATGGAGGAAATAATAT<br>GGTTTTCATTGATCTCCTTGAGTTGATTCTTAGTAATCCAGAGATCTTTGAAATTTTAC<br>TGCTGTGGGTCCACATTAACCGGGAAGACGTGGTAAAAAAGCTGTTTGATGTCATTA<br>AATGGGCAGTCGAGGATGGTGGAAACAATATGGTATTTATCCATTTGTTGCGAAAAC<br>TGTTATCCAACCCGGAAATTTTTGAAATCTTACTCTTGTGGGTCGATATCGGCAGAGA<br>AGAGGTTGTGAAAAAATTTTTTGAAAAGGTCCTGGAAGCAGTGAAAGAAGGCGGAA<br>ATGACATGATTGAAATCAAGAAGTTAGAAGAAATTCTCTTGGATCCGGAGAAATTTGA<br>GGAGTTACTTCTTGAGGTGGGCTCTTTGGAGTCCGGCGGGTCCGGTGGCTCTGGT<br>GGCAGTGATGTTTTCTTGGGTTAATTGAGGCCTGAGTATAAGGTGACTTATACTTGTA<br>ATCTATCTAAACGGGGAACCTCTCTAGTAGACAATCCCGTGCTAAATTGTAGGACTAA<br>TTCCATTTATCAGATTTCTAGAAACAGATAGGCCCTCTTCGGAGGGCCTATCTGTTTT<br>TTTTT |
| *ankyrin* | GGAATTGTGAGCGGATAACAATTCCCCTCTAGAGGGAGACCCTCGAATGGAATTGG<br>TTCTACATAAATGCCTAACGACTATCCCTTTGGGGAGTAGGGTCAAGTGACTCGAAA<br>CGATAGACAACTTGCTTTAACAAGTTGGAGATATAGTCTGCTCTGCATGGTGACATG<br>CAGCTGGATATAATTCCGGGGTAAGATTAACGACCTTATCTGAACATAATGCTACCGT<br>TTGGCTCGGGTGGATCCGGCGGATCCGGCGGTTCGGGACATCATCACCACCATCA<br>TTCTGGTATCATTCAATTAAAGGGGGTGTTAACAATGGACTACAAGGACGACGATGA<br>CAAGGATCTGGGAAAAAAGTTACTAGAAGCAGCAAGAGCTGGACAGGATGATGAGG<br>TTAGAATACTCATGGCAAACGGGGCCGATGTTAATGCAGACGATACTTGGGGATGGA<br>CCCCTCTTCATTTGGCTGCCTATCAGGGACATTTAGAGATTGTGGAAGTTCTACTTAA<br>AAATGGCGCTGACGTAAACGCTTATGACTATATTGGTTGGACTCCTTTACACTTAGCC<br>GCAGATGGTCACCTTGAAATAGTTGAAGTCTTATTAAAGAATGGAGCGGATGTCAAT<br>GCGAGTGATTACATAGGAGATACACCACTACATCTAGCTGCACATAATGGGCATCTC<br>GAAATTGTAGAGGTATTACTGAAACACGGTGCTGATGTGAATGCTCAAGATAAATTCG<br>GTAAAACAGCTTTTGATATTAGTATAGACAATGGTAACGAAGACCTAGCAGAAATCTT<br>ACAGGGCGGATCCGGCGGGTCAGGCGGGTCAGGCGATGTTTTCTTGGGTTAATTG<br>AGGCCTGAGTATAAGGTGACTTATACTTGTAATCTATCTAAACGGGGAACCTCTCTAG<br>TAGACAATCCCGTGCTAAATTGTAGGACTAATTCCATTTATCAGATTTCTAGAAACAGA<br>TAGGCCCTCTTCGGAGGGCCTATCTGTTTTTTTTT |
| *Bs prfB frameshifting element* | GACTTTAGGGGGTCTCTTTGACCTCGAA |
| *tasA*$^{ss}$ | ATGGGTATTAAAAAGAAACTCAGTCTTGGAGTTGCCTCTGCAGCACTTGGATTGGCT<br>CTCGTTGGAGGAGGAACATGGGCAGCATTTAACGACATTAAATCAAAGGATGCTACC<br>TTTGCA |
| *dsbA*$^{ss}$ | ATGGGAAAAAAGATTTGGCTGGCGCTGGCTGGTCTCGTTTCTCCGTTTAGCGCATC<br>GGCGGGAGGT |

| | |
|---|---|
| *nikA*$^{ss}$ | ATGCTCTCCACACTCCGCCGCACTCTATTTGCGCTGCTGGCTTGTGCGTCTTTTATC GTCCATGCCGCTGCACCA |
| *torT*$^{ss}$ | ATGCGCGTACTGCTATTTTTACTTCTTTCCCTTTTCATGTTGCCGGCATTTTCGGCT |
| *tasA*$^{ssnative}$ | ATGGGTATGAAAAAGAAATTGAGTTTAGGAGTTGCTTCTGCAGCACTAGGATTAGCT TTAGTTGGAGGAGGAACATGGGCAGCATTTAACGACATTAAATCAAAGGATGCTACT TTTGCA |

Note: LacO operators and T7 terminators are highlighted in gray. Td intron sequences were highlighted in cyan. Both external and internal locker sequences were colored in red while the coding sequences were colored in light blue.
